# Engineering heterothallic strains in fission yeast

**DOI:** 10.1101/2023.09.13.557521

**Authors:** Daniel García-Ruano, Ian Hsu, Baptiste Leray, Bénédicte Billard, Gianni Liti, Damien Coudreuse

## Abstract

In poor nitrogen conditions, fission yeast cells mate, undergo meiosis and form spores that are resistant to deleterious environments. Natural isolates of *Schizosaccharomyces pombe* are homothallic. This allows them to naturally switch between the two *h-* and *h+* mating types with a high frequency, thereby ensuring the presence of both mating partners in a population of cells. However, alteration of the mating type locus can abolish mating type switching or reduce it to a very low frequency. Such heterothallic strains have been isolated and are common in research laboratories due to the simplicity of their use for Mendelian genetics. In addition to the standard laboratory strains, a large collection of natural *S. pombe* isolates is now available, representing a powerful resource for investigating the genetic diversity and biology of fission yeast. However, most of these strains are homothallic, and only tedious or mutagenic strategies have been described to obtain heterothallic cells from a homothallic parent. Here, we describe a simple approach to generate heterothallic strains. It takes advantage of an alteration of the mating type locus that was previously identified in a mating type switching-deficient strain and the CRISPR-Cas9 editing tool, allowing for a one-step engineering of heterothallic cells with high efficiency.

**Take away points:**

- Conventional methods for obtaining heterothallic fission yeast strains are inefficient
- We implemented a streamlined genetic editing approach to engineer heterothallism
- All fission yeast isolates reported in Jeffares et al. 2015 can be engineered
- This method enhances the exploration of genetic diversity in wild fission yeast

## Introduction

In conditions of nutrient limitation, haploid cells of the fission yeast *Schizosaccharomyces pombe* exit the cell cycle and initiate sexual reproduction, undergoing mating, meiosis and spore formation. Spores are highly resistant to environmental perturbations and can remain dormant for long periods of time. Upon exposure to favorable growth conditions, these spores germinate and cells enter their normal vegetative division cycle. Fission yeast cells can be of the *plus* (*h+*) or *minus* (*h-*) mating type, a property that is determined by the specific allele present in the active mating type locus *mat1*, located on the right arm of chromosome II. Importantly, the two silenced loci *mat2-P* and *mat3-M*, which respectively drive the *plus* and *minus* mating type phenotypes, are critical for the process of mating-type switching (MTS). Thus, in a fully wild-type homothallic *h*^*90*^ strain, cells are capable of switching their mating type through copying the genetic information of one of the silenced loci above into the *mat1* locus. Subsequent expression of the copied gene determines the mating type of each cell (Yamada-Inagawa et al., 2007). As a result, in a population of homothallic cells, both mating types are present, allowing for sexual reproduction to occur in adverse nutritional conditions. The molecular mechanisms underlying MTS have been extensively studied and rely on a complex process involving an imprint, referred to as the *switching of mating type (smt)* signal, and recombination of the donor material of the *mat2-P* or *mat3-M* locus at *mat1*, two events that are associated with DNA replication (Arcangioli & Klar, 1991; Arcangioli & Gangloff, 2023).

While natural isolates of wild-type fission yeast cells are homothallic, the strains that are most commonly used in research laboratories are heterothallic and derived from the *h*^*–S*^ *L972* and *h*^*+N*^ *L975* genotypes initially described by Urs Leupold, with all cells in a population being either *h-* or *h+*. Alternative heterothallic mutants have also been reported that display a range of alterations in their mating-type regions (Beach & Klar, 1984; Meade & Gutz, 1976; Arcangioli & Klar, 1991; Engelke et al., 1987; Styrkársdóttir et al., 1993). Importantly, the inability of these cells to undergo MTS allows for maintaining stable mating types. This is a particularly useful property for taking advantage of fission yeast genetics. For instance, combinations of mutations can easily be obtained by crossing heterothallic strains of opposite mating types, with all asci being products of the mating between cells of different genotypes. This simple crossing approach between heterothallic strains can also be used to isolate both mating types for a given genotype, as the mating type phenotype in these strains segregates in a Mendelian fashion.

Heterothallic mutants can also be experimentally obtained from homothallic strains. One of the most commonly used protocols for this relies on the selection of randomly emerging heterothallic cells within a homothallic population. For this, homothallic cells are streaked on sporulation plates and allowed to form colonies. These plates are then exposed to iodine vapors, distinguishing between homothallic colonies, in which cells have undergone mating and sporulation (dark staining), and heterothallic colonies in which mating did not occur due the absence of MTS (light yellow staining) (Meade & Gutz, 1976). As the frequency at which loss of homothallism occurs is very low (Beach & Klar, 1984), variations of this protocol have been reported to enrich for heterothallic cells. These include exposure to mutagenic agents such as nitrous oxide and UV (Meade & Gutz, 1976) or the selection and secondary streaking of colonies in which small buds of heterothallic cells emerge at the top of a spore-containing homothallic population. In the latter, these cells can grow, taking advantage of the low nutrient uptake of the spores. Although relatively robust when applied to laboratory *h*^*90*^ strains, this approach remains time-consuming and laborious.

In addition to standard laboratory strains, the use of natural fission yeast isolates has recently emerged as a powerful tool for biological studies, taking advantage of the genetic and phenotypic diversity of these populations (Brown et al., 2011; Clément-Ziza et al., 2014; Hu et al., 2015; Jeffares et al., 2015). However, most of the isolates that have been described are homothallic, making it more difficult to engage in experimental genetic studies using these strains. Importantly, the isolation of spontaneous heterothallic cells from natural fission yeast isolates using the selection protocols described above proved to be much less efficient than what is reported using more standard laboratory strains (our unpublished data). We therefore set out to establish a simple methodology to circumvent this obstacle. Here we describe the strategy and protocol that we have implemented to successfully engineer heterothallic cells from homothallic strains using CRISPR, irrespective of their specific genetic backgrounds.

## Materials and Methods

### Yeast strains and methods

Cells were grown on YE4S plates or in EMM6S liquid medium following standard methods (Hayles & Nurse, 1992; Moreno et al., 1991). The wild type *h-Msmt-0* strain (PB1623) was provided by the team of Benoit Arcangioli (Pasteur Institute, Paris, France) and was used as a template to obtain the *Msmt0_RF* and *Psmt0_RF* repair fragments. Other standard laboratory strains used in this study were PN2 (*h*^*90*^ *L968*; wild type homothallic), PN1 (*h-L972*; wild type heterothallic, referred to as JB22 in the natural isolate collection) and PN4 (*h+ L975*; wild type heterothallic). All the natural isolates used (JB840, JB878, JB902 and JB1180) have been previously described (Jeffares et al., 2015). Note that we used a derivative of JB878 in which a hygromycin cassette was inserted between the *leu1* and *top2* loci by homologous recombination.

### Oligonucleotide sequences

All primers used in this study are listed below. Also see Fig. 1 for details.

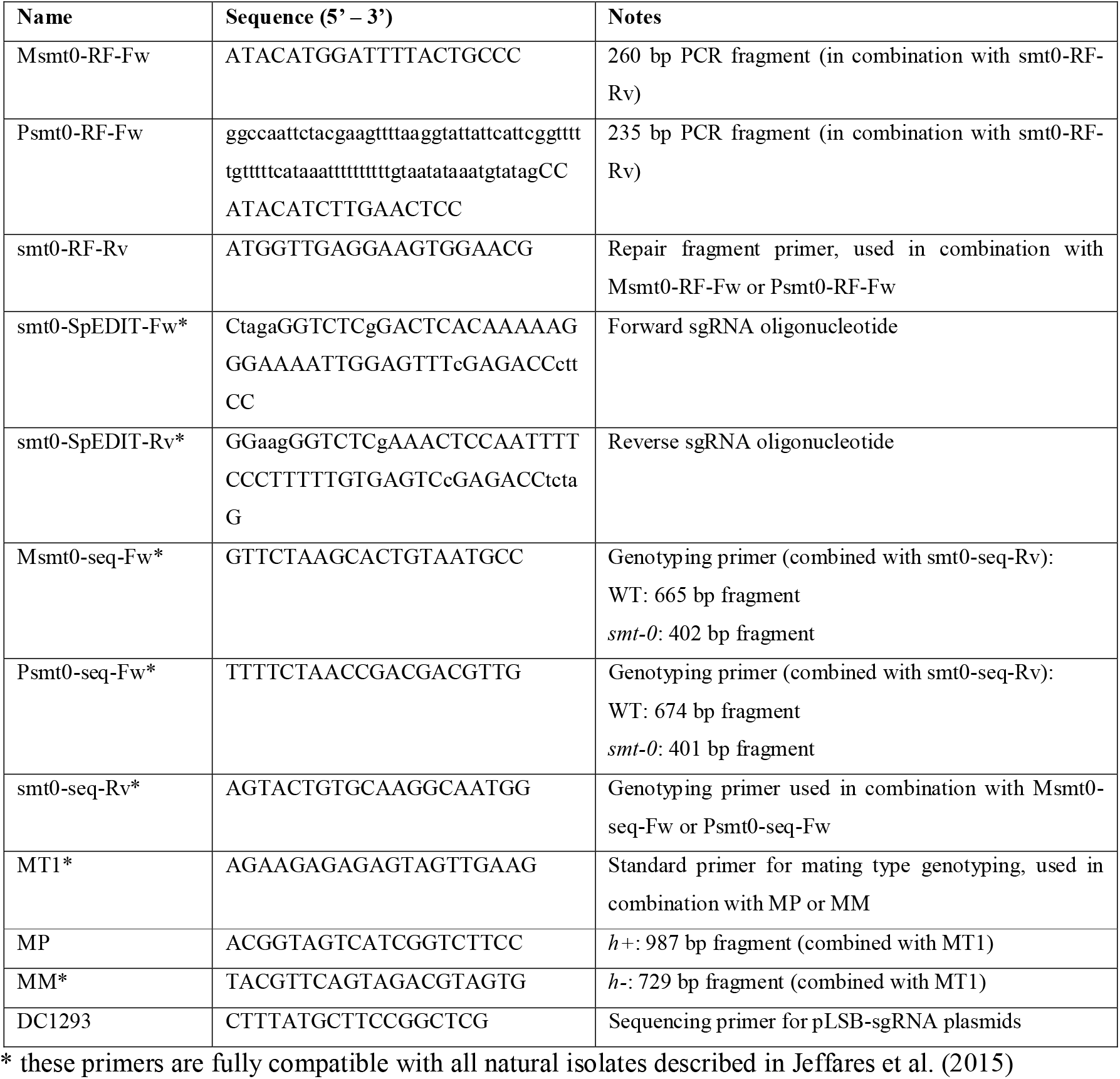

**Figure 1.**
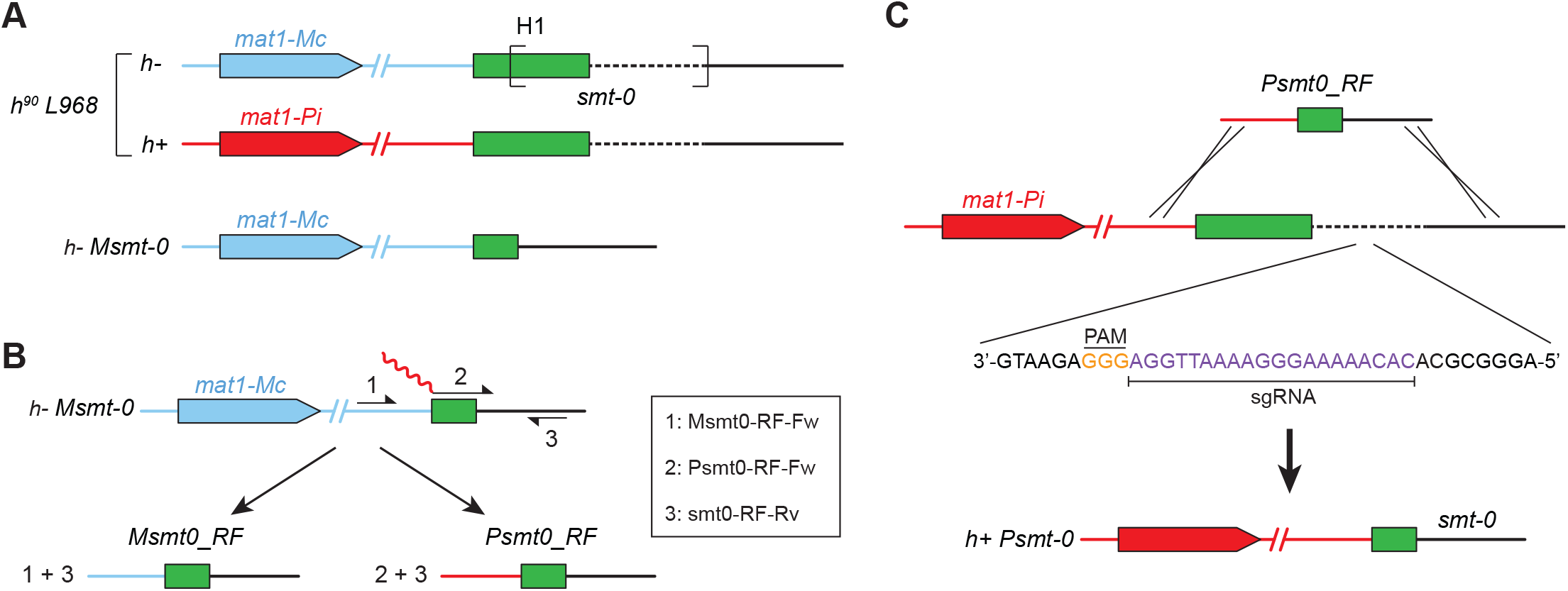
**A**. The *h*^*90*^ *L968* strain is composed of a mix of *h-* and *h+* cells that can undergo mating type switching (top panel). In the isolated heterothallic *h-Msmt-0* (bottom panel), deletion of a 263 bp region (brackets in the top panel), referred to as *smt-0*, abrogates mating type switching. **B**. Schematic of the strategy for generating repair fragments by PCR using *h-Msmt-0* DNA as a template. The combination of the Msmt0-RF-Fw and smt0-RF-Rv (1+3) amplifies the *Msmt0_RF* repair fragment. The combination of the Psmt0-RF-Fw and smt0-RF-Rv (2+3) amplifies the *Psmt0_RF* repair fragment. The 5’ part of the Psmt0-RF-Fw primer (red) corresponds to a sequence that is specific of the *mat1-P* locus of the standard laboratory strain. Note that due to the specific primer sequences that were used for generating our repair fragments, a small deletion of 9 bp (3’ end of the truncated H1 sequence, see *h-Msmt-0* in *A*) is present in the *h+ Psmt0* strains (*h-Msmt0*: ATGTATAG*TCTTTCTCC*CCATACATC; *h+ Psmt0*: ATGTATAG CCATACATC). This had no effect on the behavior of these cells but can be modified by simply altering the sequence of the Psmt0-RF-Fw primer. **C**. CRISPR-mediated editing of the mating type in homothallic cells using the repair fragments in *B* allows for fast and efficient generation of heterothallic strains. Engineering of an *h+* heterothallic strain is shown as an example. The PAM and sgRNA sequences, which are the same whether using the *Msmt0_RF* or *Psmt0_RF* repair fragments, are displayed. *B, C*: All oligonucleotide sequences are provided in the Materials and Methods section.

The editing strategy was designed according to the SpEDIT protocol (Torres-Garcia et al., 2020). Forward and reverse oligonucleotides integrating a 20 bp sequence (sgRNA inside the sequence deleted in *h-Msmt-0* cells -CACAAAAAGGGAAAATTGGA) flanked by four-base overhangs and BsaI restriction sites (smt0-SpEDIT-Fw and smt0-SpEDIT-Rv) were annealed by denaturation at 95 ºC for 30 s and subsequent cooldown to 20 ºC (∼1 ºC every 30 s). The obtained dsDNA fragment was then cloned into the pLSB-GFP-NAT plasmid using Golden Gate assembly. 10 μl of the reaction were transformed in competent *DH5a Escherichia coli* cells according to the NEB High Efficiency *E. coli* transformation protocol. Transformed cells were plated on LB-Ampicillin and incubated overnight at 37 ºC. Transformant colonies were isolated and grown to saturation in 5 mL of LB-Ampicilin medium. After spinning down the cells, all cultures showing a green pellet were discarded as the corresponding cells still contain the SpEDIT pLSB-GFP vector in which the GFP cassette has not been replaced by the sgRNA. pLSB-sgRNA plasmids were isolated from the candidate cultures (white pellets) using a standard plasmid isolation kit (NucleoSpin Plasmid, Macherey Nagel). Correct sgRNA integration was checked by sequencing using the DC1293 primer (see above).

### Amplification of the repair fragments

The repair fragments (RF) for the *smt-0* deletion were generated by PCR for both mating types. *Msmt-0_RF* was obtained using the Msmt0-RF-Fw and smt0-RF-Rv primers and genomic DNA from a *h-Msmt-0* strain. *Psmt-0_RF* was obtained similarly using the Psmt0-RF-Fw and smt0-RF-Rv primers (the Psmt0-RF-Fw primer anneals to part of the H1 sequence as well as downstream of the *smt-0* deletion and integrates a 5’ homology tail to the *mat1-P* locus – see Fig. 1). PCR reactions were purified using a PCR/Gel purification kit (NucleoSpin Gel and PCR Clean□up, Macherey-Nagel).

### Yeast electroporation

Cells were grown to exponential phase and 10^8^ cells were collected by centrifugation at 4 ºC and kept on ice throughout the rest of the procedure. A series of washing steps was then applied: 50 mL of cold (4 ºC) ultrapure water, and then 50 mL, 10 mL and 1 mL of cold 1M sorbitol. Cells were then pelleted and resuspended in 100 μL of cold 1M sorbitol. 200 ng of the pLSB-sgRNA plasmid and 1 μg of repair fragment were then added, and cells were incubated on ice for 5 min. To reduce electroporation failure, both the repair fragment and plasmid stocks were at high concentrations (around 500 and 200 ng/L, respectively) so that only small volumes (2 and 1 μL, respectively) were added to the cell suspension. Cells were then transferred to a 1 mm-gap electroporation cuvette kept on ice and electroporated using a BioRad MicroPulser Electroporator (10 μF capacitor and 600 Ω parallel resistor) configured on manual mode with a voltage of 1.5 kV. The time constant measured after the electroporation should be close to 5 ms. Cells were then immediately resuspended in 900 μL of cold 1M sorbitol, collected by centrifugation at 4 ºC, resuspended in pre-warmed YE4S medium, and incubated under shaking at 25 ºC for 3-4 h. After this recovery step, cells were divided in three aliquots of 100 μL, 900 μL and 4 mL, spun down, resuspended in 100 μL of YE4S and plated on appropriate selective medium (nourseothricin for the pLSB backbone that was used). Plates were incubated at 32 ºC for three days. Single colonies were then isolated and patched on YE4S to allow for the loss of the pLSB-sgRNA plasmid.

### *Identification of positive* transformants

Colony PCRs were performed to identify positive yeast transformants by comparing the size of the amplified fragments (see above). Different forward oligonucleotides were used for each mating type (Msmt0-seq-Fw and Psmt0-seq-Fw) in combination with a common reverse oligonucleotide (smt0-seq-Rv). To confirm the editing, the fragments were purified using a PCR/Gel purification kit (NucleoSpin Gel and PCR Clean□up, Macherey-Nagel) and sequenced using the smt0-seq-Rv oligonucleotide. The mating type of the positive candidates was further confirmed by PCR using a standard strategy, with a forward primer specific of the upstream region of the *mat1* locus (MT1), and reverse primers specific of the plus (MP) and minus (MM) mating type alleles.

### Genetic crosses

Fresh cells were patched on sporulation plates (EMM4S -N +Glutamate) either individually (test for homothallism) or together with either *h+* or *h-* heterothallic wild type cells for evaluating their mating type. Plates were then incubated at 25 ºC for three days and the presence of asci in the patch was checked by standard transmitted light microscopy.

### Microscopy

All microscopy DIC images were acquired using an inverted Zeiss Axio Observer (Carl Zeiss Microscopy GmbH, Jena, Germany) equipped with a Plan-Apochromat 63×/1.4 NA immersion lens and an Orca Flash 4.0V2 sCMOS camera (Hamamatsu Photonics, Hamamatsu, Japan). Images were acquired using the VisiView software (Visitron Systems GmbH, Puchheim, Germany).

## Results

### Design of the genome editing strategy

One of the heterothallic mutants previously isolated, referred to as the *h-Msmt-0* strain, harbors a 263 bp deletion that removes part of the H1 recombination sequence of the *mat1* locus as well as a downstream fragment (Styrkársdóttir et al., 1993) (Fig. 1A). This strain has been widely used to investigate various processes, including the mechanisms underlying MTS (Heim, 1990; Klar et al., 1991; Yamada-Inagawa et al., 2007; Bähler et al., 1993; Osman et al., 2003; Villahermosa et al., 2017; Zahedi et al., 2023). In these cells, the sequence required to induce the imprint is deleted, abolishing their capacity to undergo MTS. We therefore reasoned that this single genetic alteration could be engineered in any homothallic strain with sufficient sequence homology at the mating type locus using the CRISPR/Cas9-based SpEDIT genome editing strategy (Torres-Garcia et al., 2020).

First, we used the Benchling CRISPR guide design tool to identify potential NGG protospacer adjacent motifs (PAM) within the region deleted in *h-Msmt-0* cells. This allowed us to select a 20 bp sgRNA based on the following four criteria: (1) maximize the on-target score, (2) maximize the off-target score, (3) centered position of the sgRNA within the *smt* region, and (4) absence of reported genetic variants in known natural fission yeast isolates (Jeffares et al., 2015). Oligonucleotides for this sgRNA were then hybridized and cloned in a pLSB CRISPR plasmid following the SpEDIT protocol (see Materials and Methods).

Next, we designed two short DNA mating-type specific repair fragments (RF) that carry the *smt-0* deletion to serve as editing templates after Cas9 cleavage (Fig. 1B). The 260 bp *Msmt0_RF* was obtained by PCR amplification of genomic DNA from a *h-Msmt-0* strain, using oligonucleotides downstream of the *mat1-Mc* gene and outside of the fragment deleted in *h-Msmt-0* cells (Fig. 1B). The 235 bp *Psmt0_RF* was generated using a similar approach but with a long 100 bp oligonucleotide whose first 82 bp are homologous to the *mat1-P* locus and partial H1 sequence, while the last 3’ 18 bp anneal with a region that lies downstream of the sequence deleted in *h-Msmt-0* cells (Fig. 1B). Note that both RFs contain a 24 bp sequence homologous to H1, which is present in strains of both mating types. Thus, depending on the recombination site, both *h-Msmt-0* and *h+ Psmt-0* mutants could be theoretically obtained regardless of the RF used. We therefore designed additional primers ∼50 bp upstream and downstream of the edited region (see Materials and Methods) to genotype and sequence the engineered mating type area, with the forward oligonucleotides being specific to the mating types imposed by the RF fragments. However, our results showed that the mating types of all the strains obtained using our approach corresponded to the specific RF that was transformed (see below).

Our strategy is based on an sgRNA that is common to all the natural isolates of fission yeast described in Jeffares et al. (2015). However, a number of these strains harbor single nucleotide variants (SNV) in the regions covered by the RFs, which were designed based on the reported *smt-0* deletion and the sequence of the reference laboratory wild-type strain. While this had no incidence on the efficiency of our method (see below), the RFs can easily be customized to prevent the introduction of additional SNVs in the natural isolate backgrounds. For this, we provide the RF sequences for the different strains in Jeffares et al. (2015) (Supplementary Information). Furthermore, both the sgRNA and homology regions of the RFs can be altered to be compatible with alternative strains that may show a higher degree of genetic variation at the mating type locus. Here we report a proof-of-concept of our method that has efficiently worked in all natural isolates that we tested so far.

Altogether, this strategy generates an *smt-0* deletion in the parental homothallic strain, resulting in *h+* or *h-* heterothallic strains depending on the RF used (Fig. 1C).

### Generating heterothallic strains by engineering smt-0 in homothallic cells

As a first test, we generated heterothallic strains from the standard wild type *h*^*90*^ *L968* background. To this end, cells were co-transformed with the constructed pLSB-sgRNA plasmid and either the *Psmt0_RF* or *Msmt0_RF* repair fragment to obtain wild type *h+ Psmt-0* and *h-Msmt-0* strains, respectively. The transformation efficiency (cfu/cell number) was ∼10^-5^ for both combinations. Following the SpEDIT protocol (see Materials and Methods) and loss of the pLSB-sgRNA plasmid, a set of 28 candidates for each potential mating type was genotyped by colony PCR using the strategy described above. Remarkably, all tested candidates showed the correct *smt-0* deletion and mating type specificity. To further validate these results, we isolated single colonies for 3 candidates of each mating type, repeated the genotyping PCR, sequenced the amplified fragments, and validated the editing using alternative standard PCR reactions that allow for distinguishing between mating type alleles at the *mat1* locus (see Materials and Methods). Again, all these experiments demonstrated that our strategy allowed for editing of the mating type locus of fission yeast.

These results prompted us to assess the efficiency of our method for generating heterothallic *h-Msmt-0* and *h+ Psmt-0* strains from homothallic natural isolates of fission yeast. As a proof-of-concept, we selected JB840, JB878, JB902 as well as the sexually isolated kombucha strain JB1180 (Jeffares et al., 2015). A range of transformation efficiencies has been previously reported for fission yeast natural isolates (López Hernández et al., 2021). Using the protocol described in the Materials and Methods section, we obtained transformation efficiencies ranging from 10^-6^ to 10^-7^. While this was significantly lower than when using *h*^*90*^ *L968* cells, the correct *smt-0* deletion was found in 75-100% of the tested clones, depending on the isolate.

Collectively, this suggests that our strategy using CRISPR for mating type editing in order to generate heterothallic strains is simple, rapid, efficient and compatible with a range of genetic backgrounds, from the standard laboratory strain to natural isolates.

### Testing mating type switching in positive candidates

We next tested whether the heterothallic strains generated using our approach show the appropriate selectivity in mating. To this end, we 1) patched the engineered heterothallic stains on mating plates to assess the formation of asci, 2) crossed the candidates with *h-L972* and *h+ L975* strains to test their capacity to only mate with cells of the opposite mating type. As anticipated for *h*^*90*^ *L968* and all the natural isolates that we tested, the presence of both mating types in the populations, due to the capacity of these homothallic strains to undergo MTS, led to the formation of asci on mating plates (Fig. 2A). In contrast, the *h-Msmt-0* and *h+ Psmt-0* strains that we engineered behaved similarly to *h-L972* and *h+ L975*: only small starved cells were observed when patching these strains individually, confirming that they have lost their capacity for MTS (Fig. 2B, C). Importantly, we also found that the *h-Msmt-0* and *h+ Psmt-0* strains could only mate with *h+ L975* and *h-L972*, respectively (Fig. 2B, C). This showed specificity in mating partner, indicating that these strains are heterothallic and display the expected mating type phenotype.

**Figure 2.**
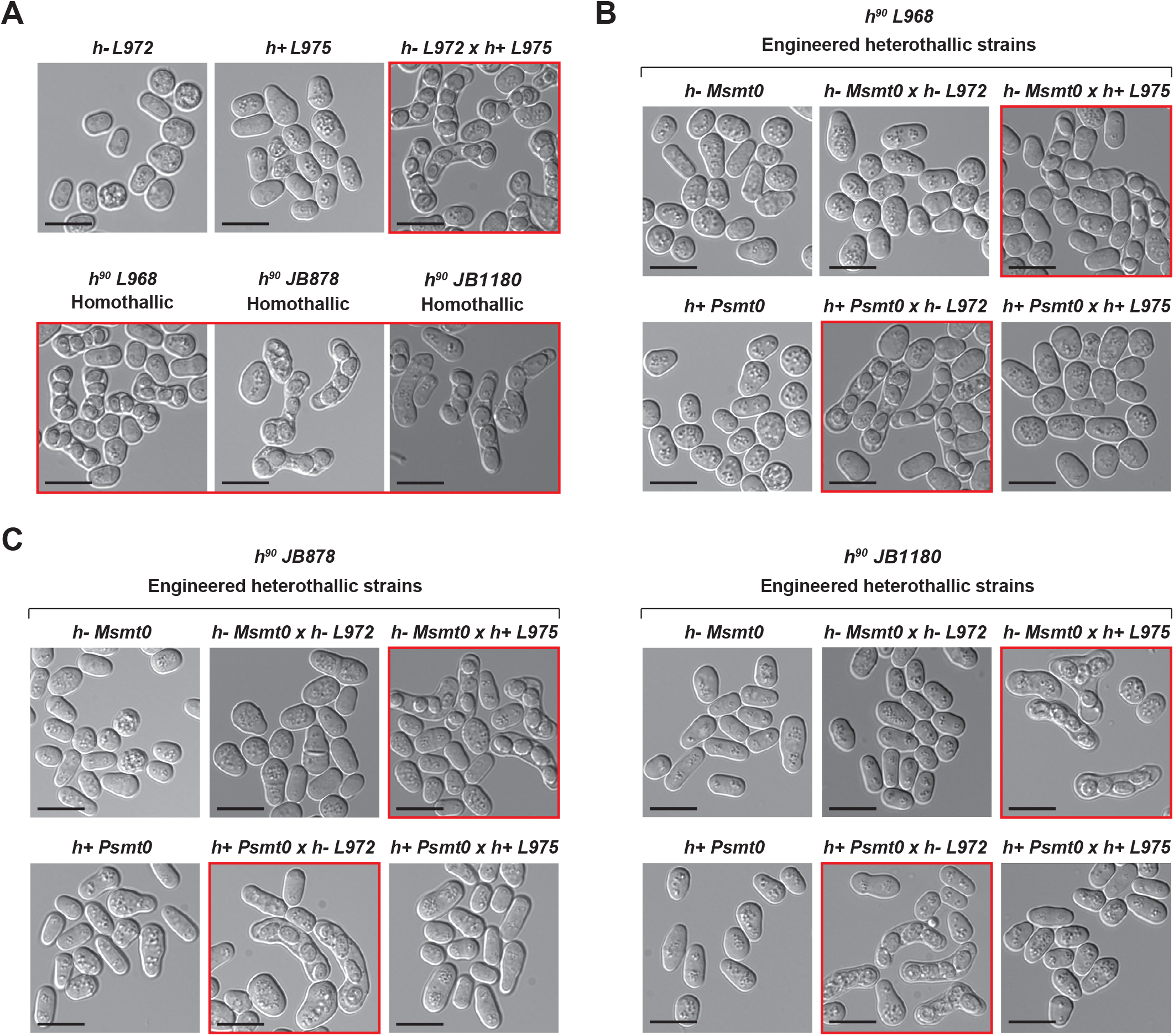
**A**. Top panel: heterothallic *h-L972* and *h+ L975* were patched on mating plates either individually or mixed (*h-L972* x *h+ L975*). The formation of asci was only detected when the two strains were crossed with each other. Bottom panel: homothallic *h*^*90*^ *L968* as well as the natural isolates *h*^*90*^ *JB878* and *h*^*90*^ *JB1180* were patched on mating plates. Due to the capacity of these cells to undergo mating type switching, asci could be observed for all three strains. **B**. *h-Msmt0* and *h+ Psmt0* heterothallic strains engineered from *h*^*90*^ *L968* were individually patched on mating plates or crossed with either *h-L972* or *h+ L975*. Asci were only observed when these strains were mixed with cells of the opposite mating type, demonstrating the effectiveness of our strategy. **C**. Assays similar to those in *B* were performed using the heterothallic strains that we engineered from the homothallic natural isolates *h*^*90*^ *JB878* and *h*^*90*^ *JB1180*. All strains that were obtained were heterothallic. Similar results were obtained using *h*^*90*^ *JB840* and *h*^*90*^ *JB902* (data not shown). *A-C*: scale bar = 10 μm. Red squares indicate conditions in which asci were observed.

## Conclusion

Our results demonstrate that targeted editing of the mating type locus using our strategy allows for generating isogenic heterothallic strains of either mating type from any homothallic population. As discussed earlier, while specific isolates may require slight adaptation of the sgRNA and RF sequences, our study suggests that the largest set of fully sequenced fission yeast natural isolates reported to date can be engineered using the tools that we describe (Jeffares et al., 2015). Our method is simple and efficient, in contrast to standard selection-based protocols or targeted strategies that involve multiple steps and low-frequency recombination events (Heim, 1990). Furthermore, while spontaneous heterothallic strains have been previously isolated, most of them carry large rearrangements of the mating type locus and many are able to revert to full homothallism (Beach & Klar, 1984). We therefore believe that our strategy is an ideal alternative to the commonly used approaches and will be particularly useful for taking full advantage of the rich biology of fission yeast natural isolates.

## Acknowledgements

We thank Pei-Yun Jenny Wu for critically reading the manuscript. This work was supported by the Agence Nationale de la Recherche (PRC eVOLve, ANR-18-CD13-0009 to D.C.), the Région Nouvelle Aquitaine (program CHESS, grant agreements 15963520 and 15964420 to D.C.), and the *Ligue contre le Cancer* (comités départementaux Gironde and Dordogne). D.G.R., I.H. and B.B. were supported by the ANR eVOLve. D.G.R. was also supported by funding from the Région Bretagne (ARED) and the Fondation ARC. B.L. was supported by the program CHESS. D.G.R., I.H., B.L. and B.B. performed the experiments. D.G.R. and D.C. designed the strategy and wrote the manuscript. All authors edited the manuscript.

