## Supplementary Information for "Engineering heterothallic strains in fission yeast"

Sequences from the start of the RF to H1 (for minus mating type)

**>JB4;JB22;JB374;JB588;JB592;JB593;JB594;JB758;JB759;JB760;JB761;JB762;JB763;JB837;JB838;JB845;JB846;JB847;JB848;JB849;JB853;JB854;JB859;JB861;JB862;JB864;JB866;JB868;JB869;JB870;JB873;JB875;JB876;JB878;JB879;JB882;JB884;JB885;JB886;JB887;JB888;JB889;JB891;JB892;JB893;JB894;JB897;JB898;JB899;JB900;JB901;JB902;JB905;JB906;JB908;JB909;JB910;JB911;JB912;JB913;JB914;JB915;JB917;JB918;JB929;JB930;JB931;JB932;JB933;JB934;JB936;JB937;JB938;JB939;JB940;JB941;JB942;JB943;JB944;JB945;JB946;JB947;JB948;JB952;JB953;JB953;JB1108;JB1109;JB1110;JB1111;JB1112;JB1113;JB1114;JB1115;JB1116;JB1117;JB1153;JB1154;JB1166;JB1167;JB1168;JB1169;JB1170;JB1171;JB1172;JB1174;JB1175;JB1176;JB1177;JB1178;JB1179;JB1180;JB1181;JB1182;JB1183;JB1184;JB1185;JB1186;JB1187;JB1188;JB1189;JB1190;JB1191;JB1192;JB1193;JB1194;JB1195;JB1196;JB1197;JB1198;JB1199;JB1201;JB1202;JB1203;JB1204;JB1205;JB1206;JB1207.chr2:2115288-2115362**

ATACATGGATTTTACTGCCCTGATTCTATCGAAATATGCTGTTTTTTTTATTCGTTTTTATTTATTTTCAATAAT

**>JB839;JB840;JB841;JB842;JB850;JB851;JB852;JB855;JB856;JB857;JB858.chr2:2115288-2115362**

ATACATGGATTTTACTGCCCTGATTCTATCGAAATATGCTGTTTTTTTTATTCGTTTTTATTTATTTTCAATATT

**>JB874.chr2:2115288-2115362**

ATACATGGATTTTACTGCCCTGATTCTATCGAAATATGCTGTTTTTTTTATTTGTTTTTATTTATTTTCAATAAT

Sequences from the start of the RF to H1 (for plus mating type)

**>JB374.MATR:4440-4498**

AGCCAATTCTACGAAGTTTTAGGGTATTATTCATTCGGTTTTTGTTTTTCATAAATTTT

**>JB758;JB866.MATR:4440-4498**

AGCCAATTCTACGAAGTTTTAGGGTATTATTCATTCGTTTTTTGTTTTTCATAAATTTC

**>JB4;JB592;JB593;JB763;JB837;JB838;JB839;JB840;JB841;JB842;JB845;JB846;JB847;JB848;JB849;JB850;JB851;JB852;JB853;JB854;JB855;JB856;JB857;JB858;JB859;JB862;JB873;JB875;JB878;JB882;JB884;JB893;JB899;JB902;JB905;JB909;JB910;JB912;JB913;JB914;JB915;JB917;JB930;JB932;JB933;JB934;JB939;JB942;JB946;JB948;JB952;JB953;JB1116;JB1153;JB1154;JB1166;JB1167;JB1171;JB1172;JB1178;JB1180;JB1185;JB1186;JB1187;JB1188;JB1196;JB1197;JB1199;JB1202;JB1203;JB1205;JB1206;JB1207.MATR:4440-4498**

AGCCAATTCTACGAAGTTTTAGGGTATTATTCATTCGTTTTTTGTTTTTCATAAATTTT

**>JB588;JB861;JB864;JB876;JB894;JB1108;JB1112;JB1113;JB1176;JB1177;JB1198;JB1201.MATR:4440-4498**

AGCCAATTCTTCGAAGTTTTAGGGTATTATTCATTCGTTTTTTGTTTTTCATAAATTTT

**>JB22;JB594;JB759;JB760;JB761;JB762;JB868;JB869;JB870;JB874;JB879;JB885;JB886;JB887;JB888;JB889;JB891;JB892;JB897;JB898;JB900;JB901;JB906;JB908;JB911;JB918;JB929;JB931;JB936;JB937;JB938;JB940;JB941;JB943;JB944;JB945;JB947;JB1109;JB1110;JB1111;JB1114;JB1115;JB1117;JB1168;JB1169;JB1170;JB1174;JB1175;JB1179;JB1181;JB1182;JB1183;JB1184;JB1189;JB1190;JB1191;JB1192;JB1193;JB1194;JB1195;JB1204.MATR:4440-4498**

GGCCAATTCTACGAAGTTTTAAGGTATTATTCATTCGGTTTTTGTTTTTCATAAATTTT

Sequences from H1 to the end of the RF (common for both mating types)

**>JB866.chr2:2115363-2115811**

TTTTTTGTAATATAAATGTATAGTCTTTCTCCTTTGTTTTCTCTCGTTCGTTTCCATGGTTTGTGATTATGCTGTTCGTGTCATTCATTCTCCCTCCAATTTTCCCTTTTTGTGTGCGCCCTTCTAACTTCCCCTTCCCTCTAACGAGCTATTTGCTTCGCTACGCTACGCACTCCCTACCATAATATACTCACTAACATTATACTTTCACTACATCTCATTACTTATTCCTACCATACACACTATATTTCAACACTTTCCTCACTCAACCATTCCATCAACCATCCTCTTTCTCACCATACATCCTGAACTCCTTACACCCATACTTCATATCCATTAGCCATTTTCCTTTATAAACATATATTAAACCTCATCCCAAATGCATCTACCCACTCTTTACATTCACATCCCTATTCCTTTAATATCTTACGTTCCACTTCCTCAACCATA

**>JB758.chr2:2115363-2115811**

TTTTTTGTAATATAAATGTATAGTCTTTCTCCTTTGTTTTCTCTCGTTCGTTTCCATGGTTTGTGATTATGCTGTTCGTGTCATTCATTCTCCCTCCAATTTTCCCTTTTTGTGTGCGCCCTTCTAACTTCCCCTTCCCTCTAACGAGCTATTTGCTTCGCTACGCTACGCACTCCCTACCATAATATACTCACTAACATTATACTTTCACTACATCTCATTACTTATTCCTACCATACACACTATATTTCAACACTTTCCTCACTCAACCATTCCATCAACCATCCTCTTTCTCACCATACATCTTGAACTCCTTACACCAATACTTCATATCCATTAGCCATTTTCCTTTATAAACATATATTAAACCTCATCCCAAATGCATCTACCCACTCTTTACATTCACATCCCTATTCCTTTAATATCTTACGTTCCACTTCCTCAACCATA

**>JB913.chr2:2115363-2115811**

TTTTTTGTAATATAAATGTATAGTCTTTCTCCTTTGTTTTCTCTCGTTCGTTTCCATGGTTTGTGATTATGCTGTTCGTGTCATTCATTCTCCCTCCAATTTTCCCTTTTTGTGTGCGCCCTTCTACCTTCCCCTTCCCTCTAACGAGATACTTGCTTCGCTACGCTACGCACTCCCTACCAAAATATACTCACTAATATTATACTTTCACTACATCTCATTACTCATTCCAACCATACACACTATATTTCAACACTTTCCTCACTCAACCACTCCATCAACCATCCTCTTTCTCCCCATACATCTTGAACTCCTTACACCAATACTTCATATCCATTAGCCATTTTCCTTTATAAACATATATTAAACCTCATCCCAAATGCATCTACCCACTCTTTACATTCACATCCCTATTCCTTTAATATCTTACATTCCACTTCCTCAACCATA

**>JB1178.chr2:2115363-2115811**

TTTTTTGTAATATAAATGTATAGTCTTTCTCCTTTGTTTTCTCTCGTTCGTTTCCATGGTTTGTGATTATGCTGTTCGTGTCATTCATTCTCCCTCCAATTTTCCCTTTTTGTGTGCGCCCTTCTACCTTCCCCTTCCCTCTAACGAGATACTTGCTTCGCTACGCTACGCACTCCCTACCAAAATATACTCACTAATATTATACTTTCACTACATCTCATTACTCATTCCAACCATACACACTATATTTCAACACTTTCCTCACTCAACCACTCCATCAACCATCCTCTTTCTCCCCATACATCTTGAACTCCTTACACCAATACTTCATATCCATTAGCCATTTTCCTTTATAAACATATATTAAACCTCATCCCAAATGCATCTACCCACTCTTTACATTCACATCCCTATTCCTTTAATATCTTACGTTCCACTTCCTCAACCATA

**>JB1172.chr2:2115363-2115811**

TTTTTTGTAATATAAATGTATAGTCTTTCTCCTTTGTTTTCTCTCGTTCGTTTCCATGGTTTGTGATTATGCTGTTCGTGTCATTCATTCTCCCTCCAATTTTCCCTTTTTGTGTGCGCCCTTCTACCTTCCCCTTCCCTCTAACGAGATACTTGCTTCGCTACGCTACGCACTCCCTACCAAAATATACTCACTAATATTATACTTTCACTACATCTCATTACTCATTCCAACCATACACACTATATTTCAACACTTTCCTCACTCACCCACTCCATCAACCATCCTCTTTCTCCCCATACATCTTGAACTCCTTACACCAATACTTCATATCCATTAGCCATTTTCCTTTATAAACATATATTAAACCTCATCCCAAATGCATCTACCCACTCTTTACATTCACATCCCTATTCCTTTAATATCTTACGTTCCACTTCCTCAACCATA

**>JB1166.chr2:2115363-2115811**

TTTTTTGTAATATAAATGTATAGTCTTTCTCCTTTGTTTTCTCTCGTTCGTTTCCATGGTTTGTGATTATGCTGTTCGTGTCATTCATTCTCCCTCCAATTTTCCCTTTTTGTGTGCGCCCTTCTACCTTCCCCTTCCCTCTAACGAGATACTTGCTTCGCTACGCTACGCACTCCCTACCAAAATATACTCACTAATATTATACTTTCACTACATCTCATTACTCATTCCAACCATACACACTATATTTCAACACTTTCCTCACTCACCCATTCCATCAACCATCCTCTTTCTCCCCATACATCTTGAACTCCTTACACCAATACTTCATATCCATTAGCCATTTTCCTTTATAAACATATATTAAACCTCATCCCAAATGCATCTACCCACTCTTTACATTCACATCCCTATTCCTTTAATATCTTACGTTCCACTTCCTCAACCATA

**>JB846;JB873;JB902.chr2:2115363-2115811**

TTTTTTGTAATATAAATGTATAGTCTTTCTCCTTTGTTTTCTCTCGTTCGTTTCCATGGTTTGTGATTATGCTGTTCGTGTCATTCATTCTCCCTCCAATTTTCCCTTTTTGTGTGCGCCCTTCTACCTTCCCCTTCCCTCTAACGAGATACTTGCTTCGCTACGCTACGCACTCCCTACCAAAATATACTCACTAATATTATACTTTCACTACATCTCATTACTCATTCCAACTATACACACTATATTTCAACACTTTCCTCACTCAACCACTCCATCAACCATCCTCTTTCTCCCCATACATCTTGAACTCCTTACACCAATACTTCATATCCATTAGCCATTTTCCTTTATAAACATATATTAAACCTCATCCCAAATGCATCTACCCACTCTTTACATTCACATCCCTATTCCTTTAATATCTTACATTCCACTTCCTCAACCATA

**>JB838;JB952.chr2:2115363-2115811**

TTTTTTGTAATATAAATGTATAGTCTTTCTCCTTTGTTTTCTCTCGTTCGTTTCCATGGTTTGTGATTATGCTGTTCGTGTCATTCATTCTCCCTCCAATTTTCCCTTTTTGTGTGCGCCCTTCTACCTTCCCCTTCCCTCTAACGAGATACTTGCTTCGCTACGCTACGCACTCCCTACCAAAATATACTCACTAATATTATACTTTCACTACATCTCATTACTCATTCCAACTATACACACTATATTTCAACACTTTCCTCACTCAACCACTCCATCAACCATCCTCTTTCTCCCCATACATCTTGAACTCCTTACACCAATACTTCATATCCATTAGCCATTTTCCTTTATAAACATATATTAAACCTCATCCCAAATGCATCTACCCACTCTTTACATTCACATCCCTATTCCTTTAATATCTTACGTTCCACTTCCTCAACCATA

**>JB932;JB934.chr2:2115363-2115811**

TTTTTTGTAATATAAATGTATAGTCTTTCTCCTTTGTTTTCTCTCGTTCGTTTCCATGGTTTGTGATTATGCTGTTCGTGTCATTCATTCTCCCTCCAATTTTCCCTTTTTGTGTGCGCCCTTCTACCTTCCCCTTCCCTCTAACGAGATACTTGCTTCGCTACGCTACGCACTCCCTACCAAAATATACTCACTAATATTATACTTTCACTACATCTCATTACTCATTCCAACTATACACACTATATTTCAACACTTTCCTCACTCAACCACTCCATCAACCATCCTCTTTCTCCCCATACATCTTGAACTCCTTACACCAATACTTCATATCCATTAGCCATTTTCCTTTATAAACATATATTAAACCTCATCCCAAATGCATCTACCCACTCTTTACATTCACATCCCTATTCCTTTAATTTCTTACATTCCACTTCCTCAACCATA

**>JB910.chr2:2115363-2115811**

TTTTTTGTAATATAAATGTATAGTCTTTCTCCTTTGTTTTCTCTCGTTCGTTTCCATGGTTTGTGATTATGCTGTTCGTGTCATTCATTCTCCCTCCAATTTTCCCTTTTTGTGTGCGCCCTTCTACCTTCCCCTTCCCTCTAACGAGATACTTGCTTCGCTACGCTACGCACTCCCTACCAAAATATACTCACTAATATTATACTTTCACTACATCTCATTACTCATTCCAACTATACACACTATATTTCAACACTTTCCTCACTCACCCACTCCATCAACCATCCTCTTTCTCCCCATACATCTTGAACTCCTTACACCAATACTTCATATCCATTAGCCATTTTCCTTTATAAACATATATTAAACCTCATCCCAAATGCATCTACCCACTCTTTACATTCACATCCCTATTCCTTTAATATCTTACGTTCCACTTCCTCAACCATA

**>JB882.chr2:2115363-2115811**

TTTTTTGTAATATAAATGTATAGTCTTTCTCCTTTGTTTTCTCTCGTTCGTTTCCATGGTTTGTGATTATGCTGTTCGTGTCATTCATTCTCCCTCCAATTTTCCCTTTTTGTGTGCGCCCTTCTACCTTCCCCTTCCCTCTAACGAGATACTTGCTTCGCTACGCTACGCACTCCCTACCAAAATATACTCACTAATATTATACTTTCACTACATCTCATTACTCATTCCAACTATACACACTATATTTCAACACTTTCCTCACTCACCCATTCCATCAACCATCCTCTTTCTCACCATACATCTTGAACTCCTTACACCAATACTTCATATCCATTAGCCATTTTCCTTTATAAACATATATTAAACCTCATCCCAAATGCATCTACCCACTCTTTACATTCACATCCCTATTCCTTTAATATCTTACGTTCCACTTCCTCAACCATA

**>JB914.chr2:2115363-2115811**

TTTTTTGTAATATAAATGTATAGTCTTTCTCCTTTGTTTTCTCTCGTTCGTTTCCATGGTTTGTGATTATGCTGTTCGTGTCATTCATTCTCCCTCCAATTTTCCCTTTTTGTGTGCGCCCTTCTACCTTCCCCTTCCCTCTAACGAGATACTTGCTTCGCTACGCTACGCACTCCCTACCAAAATATACTCACTAATATTATACTTTCACTACATCTCATTACTCATTCCAACTATACACACTATATTTCAACACTTTCCTCACTCACCTATTCCATCAACCATCCTCTTTCTCACCATACATCCTGAACTCCTTACACCCATACTTCATATCCATTAGCCATTTTCCTTTATAAACATATATTAAACCTCATCCCAAATGCATCTACCCACTCTTTACATTCACATCCCTATTCCTTTAATATCTTACATTCCACTTCCTCAACCATA

**>JB911.chr2:2115363-2115811**

TTTTTTGTAATATAAATGTATAGTCTTTCTCCTTTGTTTTCTCTCGTTCGTTTCCATGGTTTGTGATTATGCTGTTCGTGTCATTCATTCTCCCTCCAATTTTCCCTTTTTGTGTGCGCCCTTCTACCTTCCCCTTCCCTCTAACGAGATACTTGCTTCGCTACGCTACGCACTCCCTACCAAAATATACTCACTAATATTATACTTTCACTACATCTCATTACTCATTCCAACTATACACACTATATTTCAACACTTTCCTCACTCACCTATTCCATCAACCATCCTCTTTCTCACCATACATCCTGAACTCCTTACACCCATACTTCATATCCATTAGCCATTTTCCTTTATTAACATATATTAAACCTCATCCCAAATGCATCTACCCACTCTTTACATTCACATCCCTATTCCTTTAATATCTTACATTCCACTTCCTCAACCATA

**>JB762.chr2:2115363-2115811**

TTTTTTGTAATATAAATGTATAGTCTTTCTCCTTTGTTTTCTCTCGTTCGTTTCCATGGTTTGTGATTATGCTGTTCGTGTCATTCATTCTCCCTCCAATTTTCCCTTTTTGTGTGCGCCCTTCTACCTTCCCCTTCCCTCTAACGAGATACTTGCTTCGCTACGCTACGCACTCCCTACCAAAATATACTCACTAATATTATACTTTCACTACATCTCATTACTCATTCCAACTATACACACTATATTTCAACACTTTCCTCACTCACCTATTCCATCAACCATCCTCTTTCTCACCATACATCTTGAACTCCTTACACCAATACTTCATATCCATTAGCCATTTTCCTTTATAAACATATATTAAACCTCATCCCAAATGCATCTACCCACTCTTTACATTCACATCCCTATTCCTTTAATATCTTACATTCCACTTCCTCAACCATA

**>JB849;JB878.chr2:2115363-2115811**

TTTTTTGTAATATAAATGTATAGTCTTTCTCCTTTGTTTTCTCTCGTTCGTTTCCATGGTTTGTGATTATGCTGTTCGTGTCATTCATTCTCCCTCCAATTTTCCCTTTTTGTGTGCGCCCTTCTACCTTCCCCTTCCCTCTAACGAGATACTTGCTTCGCTACGCTACGCACTCCCTACCAAAATATACTCACTAATATTATACTTTCACTACATCTCATTACTCATTCCAACTATACACACTATATTTCAACACTTTCCTCACTCACCTATTCCATCAACCATCCTCTTTCTCACCATACATCTTGAACTCCTTACACCAATACTTCATATCCATTAGCCATTTTCCTTTATAAACATATATTAAACCTCATCCCAAATGCATCTACCCACTCTTTACATTCACATCCCTATTCCTTTAATATCTTACGTTCCACTTCCTCAACCATA

**>JB884;JB933.chr2:2115363-2115811**

TTTTTTGTAATATAAATGTATAGTCTTTCTCCTTTGTTTTCTCTCGTTCGTTTCCATGGTTTGTGATTATGCTGTTCGTGTCATTCATTCTCCCTCCAATTTTCCCTTTTTGTGTGCGCCCTTCTACCTTCCCCTTCCCTCTAACGAGATACTTGCTTCGCTACGCTACGCACTCCCTACCAAAATATACTCACTAATATTATACTTTCACTACATCTCATTACTCATTCCAACTATACACACTATATTTCAACACTTTCCTCACTCACCTATTCCATCAACCATCCTCTTTCTCCCCATACATCTTGAACTCCTTACACCAATACTTCATATCCATTAGCCATTTTCCTTTATAAACATATATTAAACCTCATCCCAAATGCATCTACCCACTCTTTACATTCACATCCCTATTCCTTTAATATCTTACGTTCCACTTCCTCAACCATA

**>JB845.chr2:2115363-2115811**

TTTTTTGTAATATAAATGTATAGTCTTTCTCCTTTGTTTTCTCTCGTTCGTTTCCATGGTTTGTGATTATGCTGTTCGTGTCATTCATTCTCCCTCCAATTTTCCCTTTTTGTGTGCGCCCTTCTACCTTCCCCTTCCCTCTAACGAGATACTTGCTTCGCTACGCTACGCACTCCCTACCAAAATATACTCACTAATATTATACTTTCACTACATCTCATTACTCATTCCTACCATACACACTATATTTCAACACTTTCCTCACTCAACCACTCCATCAACCATCCTCTTTCTCACCATACATCTTGAACTCCTTACACCCATACTTCATATCCATTAGCCATTTTCCTTTATTAACATATATTAAACCTCATCCCAAATGCATCTACCCACTCTTTACATTCACATCCCTATTCCTTTAATATCTTACATTCCACTTCCTCAACCATA

**>JB912;JB946;JB948;JB953;JB1116.chr2:2115363-2115811**

TTTTTTGTAATATAAATGTATAGTCTTTCTCCTTTGTTTTCTCTCGTTCGTTTCCATGGTTTGTGATTATGCTGTTCGTGTCATTCATTCTCCCTCCAATTTTCCCTTTTTGTGTGCGCCCTTCTACCTTCCCCTTCCCTCTAACGAGATACTTGCTTCGCTACGCTACGCACTCCCTACCAAAATATACTCACTAATATTATACTTTCACTACATCTCATTACTCATTCCTACCATACACACTATATTTCAACACTTTCCTCACTCAACCACTCCATCAACCATCCTCTTTCTCCCCATACATCTTGAACTCCTTACACCAATACTTCATATCCATTAGCCATTTTCCTTTATAAACATATATTAAACCTCATCCCAAATGCATCTACCCACTCTTTACATTCACATCCCTATTCCTTTAATATCTTACGTTCCACTTCCTCAACCATA

**>JB1180.chr2:2115363-2115811**

TTTTTTGTAATATAAATGTATAGTCTTTCTCCTTTGTTTTCTCTCGTTCGTTTCCATGGTTTGTGATTATGCTGTTCGTGTCATTCATTCTCCCTCCAATTTTCCCTTTTTGTGTGCGCCCTTCTACCTTCCCCTTCCCTCTAACGAGATACTTGCTTCGCTACGCTACGCACTCCCTACCAAAATATACTCACTAATATTATACTTTCACTACATCTCATTACTCATTCCTACCATACACACTATATTTCAACACTTTCCTCACTCACCCATTCCATCAACCATCCTCTTTCTCACCATACATCTTGAACTCCTTACACCAATACTTCATATCCATTAGCCATTTTCCTTTATAAACATATATTAAACCTCATCCCAAATGCATCTACCCACTCTTTACATTCACATCCCTATTCCTTTAATATCTTACGTTCCACTTCCTCAACCATA

**>JB1153.chr2:2115363-2115811**

TTTTTTGTAATATAAATGTATAGTCTTTCTCCTTTGTTTTCTCTCGTTCGTTTCCATGGTTTGTGATTATGCTGTTCGTGTCATTCATTCTCCCTCCAATTTTCCCTTTTTGTGTGCGCCCTTCTACCTTCCCCTTCCCTCTAACGAGATACTTGCTTCGCTACGCTACGCACTCCCTACCAAAATATACTCACTAATATTATACTTTCACTACATCTCATTACTCATTCCTACCATACACACTATATTTCAACACTTTCCTCACTCACCTATTCCATCAACCATCCTCTTTCTCCCCATACATCTTGAACTCCTTACACCAATACTTCATATCCATTAGCCATTTTCCTTTATAAACATATATTAAACCTCATCCCAAATGCATCTACCCACTCTTTACATTCACATCCCTATTCCTTTAATATCTTACGTTCCACTTCCTCAACCATA

**>JB1203.chr2:2115363-2115811**

TTTTTTGTAATATAAATGTATAGTCTTTCTCCTTTGTTTTCTCTCGTTCGTTTCCATGGTTTGTGATTATGCTGTTCGTGTCATTCATTCTCCCTCCAATTTTCCCTTTTTGTGTGCGCCCTTCTACCTTCCCCTTCCCTCTAACGAGATACTTGCTTCGCTACGCTACGCACTCCCTACCAAAATATACTCACTAATATTATACTTTCACTACATCTCATTACTCATTCCTACTATACACACTATATTTCAACACTTTCCTCACTCAACCACTCCATCAACCATCCTCTTTCTCACCATACATCCTGAACTCCTTACACCAATACTTCATATCCATTAGCCATTTTCCTTTATAAACATATATTAAACCTCATCCCAAATGCATCTACCCACTCTTTACATTCACATCCCTATTCCTTTAATATCTTACGTTCCACTTCCTCAACCATA

**>JB763.chr2:2115363-2115811**

TTTTTTGTAATATAAATGTATAGTCTTTCTCCTTTGTTTTCTCTCGTTCGTTTCCATGGTTTGTGATTATGCTGTTCGTGTCATTCATTCTCCCTCCAATTTTCCCTTTTTGTGTGCGCCCTTCTACCTTCCCCTTCCCTCTAACGAGATACTTGCTTCGCTACGCTACGCACTCCCTACCAAAATATACTCACTAATATTATACTTTCACTACATCTCATTACTCATTCCTACTATACACACTATATTTCAACACTTTCCTCACTCAACCACTCCATCAACCATCCTCTTTCTCACCATACATCCTGAACTCCTTACACCCATACTTCATATCCATTAGCCATTTTCCTTTATTAACATATATTAAACCTCATCCCAAATGTATCTACCCACTCTTTACATTCACATCCCTATTCCTTTAATATCTTACGTTCCACTTCCTCAACCATA

**>JB592;JB1199.chr2:2115363-2115811**

TTTTTTGTAATATAAATGTATAGTCTTTCTCCTTTGTTTTCTCTCGTTCGTTTCCATGGTTTGTGATTATGCTGTTCGTGTCATTCATTCTCCCTCCAATTTTCCCTTTTTGTGTGCGCCCTTCTACCTTCCCCTTCCCTCTAACGAGATACTTGCTTCGCTACGCTACGCACTCCCTACCAAAATATACTCACTAATATTATACTTTCACTACATCTCATTACTCATTCCTACTATACACACTATATTTCAACACTTTCCTCACTCAACCACTCCATCAACCATCCTCTTTCTCCCCATACATCTTGAACTCCTTACACCAATACTTCATATCCATTAGCCATTTTCCTTTATAAACATATATTAAACCTCATCCCAAATGCATCTACCCACTCTTTACATTCACATCCCTATTCCTTTAATATCTTACATTCCACTTCCTCAACCATA

**>JB1115.chr2:2115363-2115811**

TTTTTTGTAATATAAATGTATAGTCTTTCTCCTTTGTTTTCTCTCGTTCGTTTCCATGGTTTGTGATTATGCTGTTCGTGTCATTCATTCTCCCTCCAATTTTCCCTTTTTGTGTGCGCCCTTCTACCTTCCCCTTCCCTCTAACGAGATACTTGCTTCGCTACGCTACGCACTCCCTACCAAAATATACTCACTAATATTATACTTTCACTACATCTCATTACTCATTCCTACTATACACACTATATTTCAACACTTTCCTCACTCAACCACTCCATCAACCATCCTCTTTCTCCCCATACATCTTGAACTCCTTACACCAATACTTCATATCCATTAGCCATTTTCCTTTATAAACATATATTAAACCTCATCCCAAATGCATCTACCCACTCTTTACATTCACATCCCTATTCCTTTAATATCTTACGTTCCACTTCCTCAACCATA

**>JB854;JB939.chr2:2115363-2115811**

TTTTTTGTAATATAAATGTATAGTCTTTCTCCTTTGTTTTCTCTCGTTCGTTTCCATGGTTTGTGATTATGCTGTTCGTGTCATTCATTCTCCCTCCAATTTTCCCTTTTTGTGTGCGCCCTTCTACCTTCCCCTTCCCTCTAACGAGATACTTGCTTCGCTACGCTACGCACTCCCTACCAAAATATACTCACTAATATTATACTTTCACTACATCTCATTACTCATTCCTACTATACACACTATATTTCAACACTTTCCTCACTCACCCATTCCATCAACCATCCTCTTTCTCCCCATACATCTTGAACTCCTTACACCAATACTTCATATCCATTAGCCATTTTCCTTTATAAACATATATTAAACCTCATCCCAAATGCATCTACCCACTCTTTACATTCACATCCCTATTCCTTTAATATCTTACATTCCACTTCCTCAACCATA

**>JB847;JB848.chr2:2115363-2115811**

TTTTTTGTAATATAAATGTATAGTCTTTCTCCTTTGTTTTCTCTCGTTCGTTTCCATGGTTTGTGATTATGCTGTTCGTGTCATTCATTCTCCCTCCAATTTTCCCTTTTTGTGTGCGCCCTTCTACCTTCCCCTTCCCTCTAACGAGATACTTGCTTCGCTACGCTACGCACTCCCTACCAAAATATACTCACTAATATTATACTTTCACTACATCTCATTACTCATTCCTACTATACACACTATATTTCAACACTTTCCTCACTCACCCATTCCATCAACCATCCTCTTTCTCCCCATACATCTTGAACTCCTTACACCAATACTTCATATCCATTAGCCATTTTCCTTTATAAACATATATTAAACCTCATCCCAAATGCATCTACCCACTCTTTACATTCACATCCCTATTCCTTTAATATCTTACGTTCCACTTCCTCAACCATA

**>JB909;JB942.chr2:2115363-2115811**

TTTTTTGTAATATAAATGTATAGTCTTTCTCCTTTGTTTTCTCTCGTTCGTTTCCATGGTTTGTGATTATGCTGTTCGTGTCATTCATTCTCCCTCCAATTTTCCCTTTTTGTGTGCGCCCTTCTACCTTCCCCTTCCCTCTAACGAGATACTTGCTTCGCTACGCTACGCACTCCCTACCAAAATATACTCACTAATATTATACTTTCACTACATCTCATTACTCATTCCTACTATACACACTATATTTCAACACTTTCCTCACTCACCTATTCCATCAACCATCCTCTTTCTCCCCATACATCTTGAACTCCTTACACCAATACTTCATATCCATTAGCCATTTTCCTTTATAAACATATATTAAACCTCATCCCAAATGCATCTACCCACTCTTTACATTCACATCCCTATTCCTTTAATATCTTACATTCCACTTCCTCAACCATA

**>JB853;JB915;JB1109;JB1196;JB1202.chr2:2115363-2115811**

TTTTTTGTAATATAAATGTATAGTCTTTCTCCTTTGTTTTCTCTCGTTCGTTTCCATGGTTTGTGATTATGCTGTTCGTGTCATTCATTCTCCCTCCAATTTTCCCTTTTTGTGTGCGCCCTTCTACCTTCCCCTTCCCTCTAACGAGATACTTGCTTCGCTACGCTACGCACTCCCTACCAAAATATACTCACTAATATTATACTTTCACTACATCTCATTACTCATTCCTACTATACACACTATATTTCAACACTTTCCTCACTCACCTATTCCATCAACCATCCTCTTTCTCCCCATACATCTTGAACTCCTTACACCAATACTTCATATCCATTAGCCATTTTCCTTTATAAACATATATTAAACCTCATCCCAAATGCATCTACCCACTCTTTACATTCACATCCCTATTCCTTTAATATCTTACGTTCCACTTCCTCAACCATA

**>JB929.chr2:2115363-2115811**

TTTTTTGTAATATAAATGTATAGTCTTTCTCCTTTGTTTTCTCTCGTTCGTTTCCATGGTTTGTGATTATGCTGTTCGTGTCATTCATTCTCCCTCCAATTTTCCCTTTTTGTGTGCGCCCTTCTACCTTCCCCTTCCCTCTAACGAGATATTTGCTTCGCTACGCTACGCACTCCCTACCATAATATACTCACTAATATTATACTTTCACTACATCTCATTACTCATTCCTACCATACACACTATACTTCAACACTTTCCTCGCTCAACCACTCCATCAACCATCCTCTTTCTCCCCATACATCTTGAACTCCTTACACCAATACTTCATATCCATTAGCCATTTTCCTTTATAAACATATATTAAACCTCATCCCAAATGCATCTACCCACTCTTTACATTCACATCCCTATTCCTTTAATATCTTACGTTCCACTTCCTCAACCATA

**>JB1174.chr2:2115363-2115811**

TTTTTTGTAATATAAATGTATAGTCTTTCTCCTTTGTTTTCTCTCGTTCGTTTCCATGGTTTGTGATTATGCTGTTCGTGTCATTCATTCTCCCTCCAATTTTCCCTTTTTGTGTGCGCCCTTCTACCTTCCCCTTCCCTCTAACGAGATATTTGCTTCGCTACGCTACGCACTCCCTACCATAATATACTCACTAATATTATACTTTCACTACATCTCATTACTCATTCCTACCATACACACTATACTTCAACACTTTCCTCGCTCAACCACTCCATCAACCATCCTCTTTCTCCCTATACATCTTGAACTCCTTACACCAATACTTCATATCCATTAGCCATTTTCCTTTATAAACATATATTAAACCTCATCCCAAATGCATCTACCCACTCTTTACATTCACATCCCTATTCCTTTAATATCTTACGTTCCACTTCCTCAACCATA

**>JB588.chr2:2115363-2115811**

TTTTTTGTAATATAAATGTATAGTCTTTCTCCTTTGTTTTCTCTCGTTCGTTTCCATGGTTTGTGATTATGCTGTTCGTGTCATTCATTCTCCCTCCAATTTTCCCTTTTTGTGTGCTCCCTTCTACCTTCCCCTTCCCTCTAACGAGATACTTGCTTCGCTACGCTATGCACTCCCTACCAAAATATACTCACTAATATTATACTTTCACTACATCTCATTACTCATTCCAACCATACACACTATATTTCAACACTTTCCTCACTCACCCACTCCATCAACCATCCTCTTTCTCCCCATACATCTTGAACTCCTTACACCAATACTTCATATCCATTAGCCATTTTCCTTTATAAACATATATTAAACCTCATCCCAAATGCATCTACCCACTCTTTACATTCACATCCCTATTCCTTTAATATCTTACGTTCCACTTCCTCAACCATA

**>JB864.chr2:2115363-2115811**

TTTTTTGTAATATAAATGTATAGTCTTTCTCCTTTGTTTTCTCTCGTTCGTTTCCATGGTTTGTGATTATGCTGTTCGTGTCATTCATTCTCCCTCCAATTTTCCCTTTTTGTGTGCTCCCTTCTACCTTCCCCTTCCCTCTAACGAGATACTTGCTTCGCTACGCTATGCACTCCCTACCAAAATATACTCACTAATATTATACTTTCACTACATCTCATTACTCATTCCAACTATACACACTATATTTCAACACTTTCCTCACTCACCCATTCCATCAACCATCCTCTTTCTCACCATACATCCTGAACTCCTTACACCAATACTTCATATCCATTAGCCATTTTCCTTTATAAACATATATTAAACCTCATCCCAAATGCATCTACCCACTCTTTACATTCACATCCCTATTCCTTTAATATCTTACGTTCCACTTCCTCAACCATA

**>JB876;JB1177.chr2:2115363-2115811**

TTTTTTGTAATATAAATGTATAGTCTTTCTCCTTTGTTTTCTCTCGTTCGTTTCCATGGTTTGTGATTATGCTGTTCGTGTCATTCATTCTCCCTCCAATTTTCCCTTTTTGTGTGCTCCCTTCTACCTTCCCCTTCCCTCTAACGAGATACTTGCTTCGCTACGCTATGCACTCCCTACCAAAATATACTCACTAATATTATACTTTCACTACATCTCATTACTCATTCCAACTATACACACTATATTTCAACACTTTCCTCACTCACCCATTCCATCAACCATCCTCTTTCTCACCATACATCTTGAACTCCTTACACCAATACTTCATATCCATTAGCCATTTTCCTTTATAAACATATATTAAACCTCATCCCAAATGCATCTACCCACTCTTTACATTCACATCCCTATTCCTTTAATATCTTACGTTCCACTTCCTCAACCATA

**>JB894.chr2:2115363-2115811**

TTTTTTGTAATATAAATGTATAGTCTTTCTCCTTTGTTTTCTCTCGTTCGTTTCCATGGTTTGTGATTATGCTGTTCGTGTCATTCATTCTCCCTCCAATTTTCCCTTTTTGTGTGCTCCCTTCTACCTTCCCCTTCCCTCTAACGAGATACTTGCTTCGCTACGCTATGCACTCCCTACCAAAATATACTCACTAATATTATACTTTCACTACATCTCATTACTCATTCCAACTATACACACTATATTTCAACACTTTCCTCACTCACCCATTCCATCAACCATCCTCTTTCTCCCCATACATCTTGAACTCCTTACACCAATACTTCATATCCATTAGCCATTTTCCTTTATAAACATATATTAAACCTCATCCCAAATGCATCTACCCACTCTTTACATTCACATCCCTATTCCTTTAATATCTTACATTCCACTTCCTCAACCATA

**>JB1113.chr2:2115363-2115811**

TTTTTTGTAATATAAATGTATAGTCTTTCTCCTTTGTTTTCTCTCGTTCGTTTCCATGGTTTGTGATTATGCTGTTCGTGTCATTCATTCTCCCTCCAATTTTCCCTTTTTGTGTGCTCCCTTCTACCTTCCCCTTCCCTCTAACGAGATACTTGCTTCGCTACGCTATGCACTCCCTACCAAAATATACTCACTAATATTATACTTTCACTACATCTCATTACTCATTCCTACCATACACACTATATTTCAACACTTTCCTCACTCAACCACTCCATCAACCATCCTCTTTCTCCCCATACATCTTGAACTCCTTACACCAATACTTCATATCCATTAGCCATTTTCCTTTATAAACATATATTAAACCTCATCCCAAATGCATCTACCCACTCTTTACATTCACATCCCTATTCCTTTAATATCTTACGTTCCACTTCCTCAACCATA

**>JB1108;JB1112;JB1198;JB1201.chr2:2115363-2115811**

TTTTTTGTAATATAAATGTATAGTCTTTCTCCTTTGTTTTCTCTCGTTCGTTTCCATGGTTTGTGATTATGCTGTTCGTGTCATTCATTCTCCCTCCAATTTTCCCTTTTTGTGTGCTCCCTTCTACCTTCCCCTTCCCTCTAACGAGATACTTGCTTCGCTACGCTATGCACTCCCTACCAAAATATACTCACTAATATTATACTTTCACTACATCTCATTACTCATTCCTACCATACACACTATATTTCAACACTTTCCTCACTCACCCATTCCATCAACCATCCTCTTTCTCCCCATACATCTTGAACTCCTTACACCAATACTTCATATCCATTAGCCATTTTCCTTTATAAACATATATTAAACCTCATCCCAAATGCATCTACCCACTCTTTACATTCACATCCCTATTCCTTTAATATCTTACGTTCCACTTCCTCAACCATA

**>JB861.chr2:2115363-2115811**

TTTTTTGTAATATAAATGTATAGTCTTTCTCCTTTGTTTTCTCTCGTTCGTTTCCATGGTTTGTGATTATGCTGTTCGTGTCATTCATTCTCCCTCCAATTTTCCCTTTTTGTGTGCTCCCTTCTACCTTCCCCTTCCCTCTAACGAGATACTTGCTTCGCTACGCTATGCACTCCCTACCAAAATATACTCACTAATATTATACTTTCACTACATCTCATTACTCATTCCTACTATACACACTATATTTCAACACTTTCCTCACTCACCCATTCCATCAACCATCCTCTTTCTCCCCATACATCCTGAACTCCTTACACCCATACTTCATATCCATTAGCCATTTTCCTTTATAAACATATATTAAACCTCATCCCAAATGCATCTACCCACTCTTTACATTCACATCCCTATTCCTTTAATATCTTACGTTCCACTTCCTCAACCATA

**>JB1176.chr2:2115363-2115811**

TTTTTTGTAATATAAATGTATAGTCTTTCTCCTTTGTTTTCTCTCGTTCGTTTCCATGGTTTGTGATTATGCTGTTCGTGTCATTCATTCTCCCTCCAATTTTCCCTTTTTGTGTGCTCCCTTCTACCTTCCCCTTCCCTCTAACGAGATACTTGCTTCGCTACGCTATGCACTCCCTACCAAAATATACTCACTAATATTATACTTTCACTACATCTCATTACTCATTCCTACTATACACACTATATTTCAACACTTTCCTCACTCACCCATTCCATCAACCATCCTCTTTCTCCCCATACATCTTGAACTCCTTACACCAATACTTCATATCCATTAGCCATTTTCCTTTATAAACATATATTAAACCTCATCCCAAATGCATCTACCCACTCTTTACATTCACATCCCTATTCCTTTAATATCTTACGTTCCACTTCCTCAACCATA

**>JB862.chr2:2115363-2115811**

TTTTTTGTAATATAAATGTATAGTCTTTCTCCTTTGTTTTCTCTCGTTCGTTTCCATGGTTTGTGATTATGCTGTTCGTGTCATTCATTCTCCCTCCAATTTTCCCTTTTTGTGTGTGCCCTTCTACCTTCCCCTTCCCTCTACCGAGATACTTGCTTCGCTACGCTACGCACTCCCTACCAAAATATACTCACTAATATTATACTTTCACTACATCTCATTACTCATTCCAACTATACACACTATATTTCAACACTTTCCTCACTCACCCATTCCATCAACCATCCTCTTTCTCCCCATACATCTTGAACTCCTTACACCAATACTTCATATCCATTAGCCATTTTCCTTTATAAACATATATTAAACCTCATCCCAAATGCATCTACCCACTCTTTACATTCACATCCCTATTCCTTTAATATCTTACATTCCACTTCCTCAACCATA

**>JB930.chr2:2115363-2115811**

TTTTTTGTAATATAAATGTATAGTCTTTCTCCTTTGTTTTCTCTCGTTCGTTTCCATGGTTTGTGATTATGCTGTTCGTGTCATTCATTCTCCCTCCAATTTTCCCTTTTTGTGTGTTCCCTTCTACCTTCCCCTTCCCTCTACCGAGATACTTGCTTCGCTACGCTACGCACTCCCTACCAAAATATACTCACTAATATTATACTTTCACTACATCTCATTACTCATTCCTACCATACACACTATATTTCAACACTTTCCTCACTCACCCATTCCATCAACCATCCTCTTTCTCACCATACATCTTGAACTCCTTACACCAATACTTCATATCCATTAGCCATTTTCCTTTATAAACATATATTAAACCTCATCCCAAATGCATCTACCCACTCTTTACATTCACATCCCTATTCCTTTAATTTCTTACGTTCCACTTCCTCAACCATA

**>JB943.chr2:2115363-2115811**

TTTTTTGTAATATAAATGTATAGTCTTTCTCCTTTGTTTTCTCTCGTTCGTTTCCATGGTTTGTGATTATGCTGTTCGTGTCATTCATTCTCCCTCCAATTTTCCCTTTTTGTGTGTTCCCTTCTACCTTCCCCTTCCCTCTACCGAGATACTTGCTTCGCTACGCTACGCACTCCCTACCAAAATATACTCACTAATATTATACTTTCACTACATCTCATTACTCATTCCTACTATACACACTATATTTCAACACTTTCCTCACTCACCCATTCCATCAACCATCCTCTTTCTCCCCATACATCTTGAACTCCTTACACCAATACTTCATATCCATTAGCCATTTTCCTTTATAAACATATATTAAACCTCATCCCAAATGCATCTACCCACTCTTTACATTCACATCCCTATTCCTTTAATATCTTACGTTCCACTTCCTCAACCATA

**>JB842.chr2:2115363-2115811**

TTTTTTGTAATATAAATGTATAGTCTTTCTCCTTTGTTTTCTCTCGTTCGTTTCCATGTTGTCCAATTATGCTGTTCGTGTCATTCATTCTCCCTCCAATTTTCCCTTTTTGTGTGCGCCCTTCTACCTTCCCCTTCCCTCTAACGAGATACTTGCTTCGCTACGCTACGCACTCCCTACCAAAATATACTCACTAATATTATACTTTCACTACATCTCATTACTCATTCCAACTATACACACTATATTTCAACACTTTCCTCACTCACCCATTCCATCAACCATCCTCTTTCTCCCCATACATCCTGAACTCCTTACACCAATACTTCATATCCATTAGCCATTTTCCTTTATAAACATATATTAAACCTCATCCCAAATGCATCTACCCACTCTTTACATTCACATCCCTATTCCTTTAATATCTTACGTTCCACTTCCTCAACCATA

**>JB850.chr2:2115363-2115811**

TTTTTTGTAATATAAATGTATAGTCTTTCTCCTTTGTTTTCTCTCGTTCGTTTCCATGTTGTCCAATTATGCTGTTCGTGTCATTCATTCTCCCTCCAATTTTCCCTTTTTGTGTGCGCCCTTCTACCTTCCCCTTCCCTCTAACGAGATACTTGCTTCGCTACGCTACGCACTCCCTACCAAAATATACTCACTAATATTATACTTTCACTACATCTCATTACTCATTCCAACTATACACACTATATTTCAACACTTTCCTCACTCACCCATTCCATCAACCATCCTCTTTCTCCCCATACATCTTGAACTCCTTACACCAATACTTCATATCCATTAGCCATTTTCCTTTATAAACATATATTAAACCTCATCCCAAATGCATCTACCCACTCTTTACATTCACATCCCTATTCCTTTAATATCTTACGTTCCACTTCCTCAACCATA

**>JB856.chr2:2115363-2115811**

TTTTTTGTAATATAAATGTATAGTCTTTCTCCTTTGTTTTCTCTCGTTCGTTTCCATGTTGTCCAATTATGCTGTTCGTGTCATTCATTCTCCCTCCAATTTTCCCTTTTTGTGTGCGCCCTTCTACCTTCCCCTTCCCTCTAACGAGATACTTGCTTCGCTACGCTACGCACTCCCTACCAAAATATACTCACTAATATTATACTTTCACTACATCTCATTACTCATTCCTACCATACACACTATATTTCAACACTTTCCTCACTCACCCATTCCATCAACCATCCTCTTTCTCACCATACATCCTGAACTCCTTACACCAATACTTCATATCCATTAGCCATTTTCCTTTATAAACATATATTAAACCTCATCCCAAATGCATCTACCCACTCTTTACATTCACATCCCTATTCCTTTAATATCTTACGTTCCACTTCCTCAACCATA

**>JB840;JB851.chr2:2115363-2115811**

TTTTTTGTAATATAAATGTATAGTCTTTCTCCTTTGTTTTCTCTCGTTCGTTTCCATGTTGTCCAATTATGCTGTTCGTGTCATTCATTCTCCCTCCAATTTTCCCTTTTTGTGTGCGCCCTTCTACCTTCCCCTTCCCTCTAACGAGATACTTGCTTCGCTACGCTACGCACTCCCTACCAAAATATACTCACTAATATTATACTTTCACTACATCTCATTACTCATTCCTACCATACACACTATATTTCAACACTTTCCTCACTCACCCATTCCATCAACCATCCTCTTTCTCCCCATACATCTTGAACTCCTTACACCAATACTTCATATCCATTAGCCATTTTCCTTTATAAACATATATTAAACCTCATCCCAAATGCATCTACCCACTCTTTACATTCACATCCCTATTCCTTTAATATCTTACGTTCCACTTCCTCAACCATA

**>JB855.chr2:2115363-2115811**

TTTTTTGTAATATAAATGTATAGTCTTTCTCCTTTGTTTTCTCTCGTTCGTTTCCATGTTGTCCAATTATGCTGTTCGTGTCATTCATTCTCCCTCCAATTTTCCCTTTTTGTGTGCGCCCTTCTACCTTCCCCTTCCCTCTAACGAGATACTTGCTTCGCTACGCTACGCACTCCCTACCAAAATATACTCACTAATATTATACTTTCACTACATCTCATTACTCATTCCTACTATACACACTATATTTCAACACTTTCCTCACTCACCCACTCCATCAACCATCCTCTTTCTCCCCATACATCTTGAACTCCTTACACCAATACTTCATATCCATTAGCCATTTTCCTTTATAAACATATATTAAACCTCATCCCAAATGCATCTACCCACTCTTTACATTCACATCCCTATTCCTTTAATATCTTACGTTCCACTTCCTCAACCATA

**>JB857.chr2:2115363-2115811**

TTTTTTGTAATATAAATGTATAGTCTTTCTCCTTTGTTTTCTCTCGTTCGTTTCCATGTTGTCCAATTATGCTGTTCGTGTCATTCATTCTCCCTCCAATTTTCCCTTTTTGTGTGCGCCCTTCTACCTTCCCCTTCCCTCTAACGAGATACTTGCTTCGCTACGCTACGCACTCCCTACCAAAATATACTCACTAATATTATACTTTCACTACATCTCATTACTCATTCCTACTATACACACTATATTTCAACACTTTCCTCACTCACCCATTCCATCAACCATCCTCTTTCTCACCATACATCTTGAACTCCTTACACCAATACTTCATATCCATTAGCCATTTTCCTTTATAAACATATATTAAACCTCATCCCAAATGCATCTACCCACTCTTTACATTCACATCCCTATTCCTTTAATATCTTACGTTCCACTTCCTCAACCATA

**>JB839;JB841;JB852;JB858.chr2:2115363-2115811**

TTTTTTGTAATATAAATGTATAGTCTTTCTCCTTTGTTTTCTCTCGTTCGTTTCCATGTTTCCAAATATGCTGTTCGTGTCATTCATTCTCCCTCCAATTTTCCCTTTTTGTGTGCGCCCTTCTACCTTCCCCTTCCCTCTAACGAGATATTTGCTTCGCTACGCTACGCACTCCCTACCATAATATACTCACTAATATTATACTTTCACTACATCTCATTACTCATTCCTACCATACACACTATACTTCAACACTTTCCTCGCTCAACCACTCCATCAACCATCCTCTTTCTCCCCATACATCTTGAACTCCTTACACCAATACTTCATATCCATTAGCCATTTTCCTTTATAAACATATATTAAACCTCATCCCAAATGCATCTACCCACTCTTTACATTCACATCCCTATTCCTTTAATATCTTACGTTCCACTTCCTCAACCATA

**>JB1188.chr2:2115363-2115811**

TTTTTTGTAATATAAATGTATAGTCTTTCTCCTTTGTTTTCTCTCGTTCGTTTCCATGTTTCCAATTATGCTGTTCGTGTCATTCATTCTCCCTCCAATTTTCCCTTTTTGTGTGCGCCCTTCTAACTTCCCCTTCCCTCTAACGAGAAATTTGCTTCGCTACGCTACGCACTCCCTACCATAATATACTCACTAACATTATACTTTCACTACATCTCATTACTTATTCCTACCATACACACTATATTTCAACACTTTCCTCACTCAACCATTCCATCAACCATCCTCTTTCTCCCCATACATCTTGAACTCCTTACACCAATACTTCATATCCATTAGCCATTTTCCTTTATAAACATATATTAAACCTCATCCCAAATGCATCTACCCACTCTTTACATTCACATCCCTATTCCTTTAATATCTTACGTTCCACTTCCTCAACCATA

**>JB899.chr2:2115363-2115811**

TTTTTTGTAATATAAATGTATAGTCTTTCTCCTTTGTTTTCTCTCGTTCGTTTCCATGTTTCCAATTATGCTGTTCGTGTCATTCATTCTCCCTCCAATTTTCCCTTTTTGTGTGCGCCCTTCTAACTTCCCCTTCCCTCTAACGAGAAATTTGCTTCGCTACGCTACGCACTCCCTACCATAATATACTCACTAACATTATACTTTCACTACATCTCATTACTTATTCCTACCATACACACTATATTTCAACACTTTCCTCACTCAACCATTCCATCAACCATCCTCTTTCTCCCCATACATCTTGAACTCCTTACACCCATACTTCATATCCATTACCCATTTTCCTTTATTAACATATATTAAACCTCATCCCAAATGCATCTACCCACTCTTTACATTCACATCCCTATTCCTTTAATATCTTACGTTCCACTTCCTCAACCATA

**>JB1186.chr2:2115363-2115811**

TTTTTTGTAATATAAATGTATAGTCTTTCTCCTTTGTTTTCTCTCGTTCGTTTCCATGTTTCCAATTATGCTGTTCGTGTCATTCATTCTCCCTCCAATTTTCCCTTTTTGTGTGCGCCCTTCTAACTTCCCCTTCCCTCTAACGAGAAATTTGCTTCGCTACGCTACGCACTCCCTACCATAATATACTCACTAACATTATACTTTCACTACATCTCATTACTTATTCCTACCATACACACTATATTTCAACACTTTCCTCGCTCAACCACTCCATCAACCATCCTCTTTCTCCCCATACATCTTGAACTCCTTACACCAATACTTCATATCCATTACCCATTTTCCTTTATTAACATATATTAAACCTCATCCCAAATGTATCTACCCACTCTTTACATTCACATCCTTATTCCTTTAATTTCTTACCTTCCACTTCCTCAACCATA

**>JB1187.chr2:2115363-2115811**

TTTTTTGTAATATAAATGTATAGTCTTTCTCCTTTGTTTTCTCTCGTTCGTTTCCATGTTTCCAATTATGCTGTTCGTGTCATTCATTCTCCCTCCAATTTTCCCTTTTTGTGTGCGCCCTTCTAACTTCCCCTTCCCTCTAACGAGAAATTTGCTTCGCTACGCTACGCACTCCCTACCATAATATACTCACTAACATTATACTTTCACTACATCTCATTACTTATTCCTACCATACACACTATATTTCAACACTTTCCTCGCTCAACCACTCCATCAACCATCCTCTTTCTCCCCATACATCTTGAACTCCTTACACCAATACTTCATATCCATTAGCCATTTTCCTTTATAAACATATATTAAACCTCATCCCAAATGCATCTACCCACTCTTTACATTCACATCCCTATTCCTTTAATATCTTACGTTCCACTTCCTCAACCATA

**>JB1185.chr2:2115363-2115811**

TTTTTTGTAATATAAATGTATAGTCTTTCTCCTTTGTTTTCTCTCGTTCGTTTCCATGTTTCCAATTATGCTGTTCGTGTCATTCATTCTCCCTCCAATTTTCCCTTTTTGTGTGCGCCCTTCTAACTTCCCCTTCCCTCTAACGAGAAATTTGCTTCGCTACGCTACGCACTCCCTACCATAATATACTCACTAACATTATACTTTCACTACATCTCATTACTTATTCCTACCATACACACTATATTTCAACACTTTCCTCGCTCAACCACTCCATCAACCATCCTCTTTCTCCCCATACATCTTGAACTCCTTACACCAATACTTCATATCCATTAGCCATTTTCCTTTATAAACATATATTAAACCTCATCCCAAATGTATCTACCCACTCTTTACATTCACATCCTTATTCCTTTAATTTCTTACCTTCCACTTCCTCAACCATA

**>JB1205;JB1206.chr2:2115363-2115811**

TTTTTTGTAATATAAATGTATAGTCTTTCTCCTTTGTTTTCTCTCGTTCGTTTCCATGTTTCCAATTATGCTGTTCGTGTCATTCATTCTCCCTCCAATTTTCCCTTTTTGTGTGCGCCCTTCTAACTTCCCCTTCCCTCTAACGAGATATTTGCTTCGCTACGCTACGCACTCCCTACCATAATATACTCACTAACATTATACTTTCACTACATCTCATTACTCATTCCTACCATACACACTATATTTCAACACTTTCCTCACTCAACCATTCCATCAACCATCCTCTTTCTCACCATACATCCTAAACTCCTTACACCCATACTTCATATCCATTACCCATTTTCCTTTATTAACATATATTAAACCTCATCCCAAATGTATCTACCCACTCTTTACATTCACATCCTTATTCCTTTAATTTCTTACCTTCCACTTCCTCAACCATA

**>JB1207.chr2:2115363-2115811**

TTTTTTGTAATATAAATGTATAGTCTTTCTCCTTTGTTTTCTCTCGTTCGTTTCCATGTTTCCAATTATGCTGTTCGTGTCATTCATTCTCCCTCCAATTTTCCCTTTTTGTGTGCGCCCTTCTAACTTCCCCTTCCCTCTAACGAGATATTTGCTTCGCTACGCTACGCACTCCCTACCATAATATACTCACTAACATTATACTTTCACTACATCTCATTACTCATTCCTACCATACACACTATATTTCAACACTTTCCTCACTCAACCATTCCATCAACCATCCTCTTTCTCACCATACATCCTGAACTCCTTACACCCATACTTCATATCCATTACCCATTTTCCTTTATTAACATATATTAAACCTCATCCCAAATGTATCTACCCACTCTTTACATTCACATCCTTATTCCTTTAATTTCTTACCTTCCACTTCCTCAACCATA

**>JB859.chr2:2115363-2115811**

TTTTTTGTAATATAAATGTATAGTCTTTCTCCTTTGTTTTCTCTCGTTCGTTTCCATGTTTCCAATTATGCTGTTCGTGTCATTCATTCTCCCTCCAATTTTCCCTTTTTGTGTGCGCCCTTCTAACTTCCCCTTCCCTCTAACGAGATATTTGCTTCGCTACGCTACGCACTCCCTACCATAATATACTCACTAACATTATACTTTCACTACATCTCATTACTTATTCCTACCATACACACTATATTTCAACACTTTCCTCACTCAACCACTCCATCAACCATCCTCTTTCTCCCCATACATCCTAAACTCCTTACACCCATACTTCATATCCATTACCCATTTTCCTTTATTAACATATATTAAACCTCATCCCAAATGTATCTACCCACTCTTTACATTCACATCCTTATTCCTTTAATTTCTTACCTTCCACTTCCTCAACCATA

**>JB1167.chr2:2115363-2115811**

TTTTTTGTAATATAAATGTATAGTCTTTCTCCTTTGTTTTCTCTCGTTCGTTTCCATGTTTCCAATTATGCTGTTCGTGTCATTCATTCTCCCTCCAATTTTCCCTTTTTGTGTGCGCCCTTCTAACTTCCCCTTCCCTCTAACGAGATATTTGCTTCGCTACGCTACGCACTCCCTACCATAATATACTCACTAACATTATACTTTCACTACATCTCATTACTTATTCCTACCATACACACTATATTTCAACACTTTCCTCACTCAACCACTCCATCAACCATCCTCTTTCTCCCCATACATCTTGAACTCCTTACACCAATACTTCATATCCATTAGCCATTTTCCTTTATAAACATATATTAAACCTCATCCCAAATGCATCTACCCACTCTTTACATTCACATCCCTATTCCTTTAATATCTTACGTTCCACTTCCTCAACCATA

**>JB1110;JB1154.chr2:2115363-2115811**

TTTTTTGTAATATAAATGTATAGTCTTTCTCCTTTGTTTTCTCTCGTTCGTTTCCATGTTTCCAATTATGCTGTTCGTGTCATTCATTCTCCCTCCAATTTTCCCTTTTTGTGTGCGCCCTTCTAACTTCCCCTTCCCTCTAACGAGATATTTGCTTCGCTACGCTACGCACTCCCTACCATAATATACTCACTAACATTATACTTTCACTACATCTCATTACTTATTCCTACCATACACACTATATTTCAACACTTTCCTCACTCAACCACTCCATCAACCATCCTCTTTCTCCCCATACATCTTGAACTCCTTACACCCATACTTCATATCCATTACCCATTTTCCTTTATTAACATATATTAAACCTCATCCCAAATGTATCTACCCACTCTTTACATTCACATCCTTATTCCTTTAATTTCTTACCTTCCACTTCCTCAACCATA

**>JB893.chr2:2115363-2115811**

TTTTTTGTAATATAAATGTATAGTCTTTCTCCTTTGTTTTCTCTCGTTCGTTTCCATGTTTCCAATTATGCTGTTCGTGTCATTCATTCTCCCTCCAATTTTCCCTTTTTGTGTGCGCCCTTCTAACTTCCCCTTCCCTCTAACGAGATATTTGCTTCGCTACGCTACGCACTCCCTACCATAATATACTCACTAACATTATACTTTCACTACATCTCATTACTTATTCCTACCATACACACTATATTTCAACACTTTCCTCACTCAACCATTCCATCAACCATCCTCTTTCTCACCATACATCTTGAACTCCTTACACCCATACTTCATATCCATTACCCATTTTCCTTTATTAACATATATTAAACCTCATCCCAAATGTATCTACCCACTCTTTACATTCACATCCTTATTCCTTTAATTTCTTACCTTCCACTTCCTCAACCATA

**>JB593.chr2:2115363-2115811**

TTTTTTGTAATATAAATGTATAGTCTTTCTCCTTTGTTTTCTCTCGTTCGTTTCCATGTTTCCAATTATGCTGTTCGTGTCATTCATTCTCCCTCCAATTTTCCCTTTTTGTGTGCGCCCTTCTAACTTCCCCTTCCCTCTAACGAGATATTTGCTTCGCTACGCTACGCACTCCCTACCATAATATACTCACTAACATTATACTTTCACTACATCTCATTACTTATTCCTACCATACACACTATATTTCAACACTTTCCTCACTCAACCATTCCATCAACCATCCTCTTTCTCCCCATACATCTTGAACTCCTTACACCAATACTTCATATCCATTACCCATTTTCCTTTATTAACATATATTAAACCTCATCCCAAATGTATCTACCCACTCTTTACATTCACATCCTTATTCCTTTAATTTCTTACCTTCCACTTCCTCAACCATA

**>JB905.chr2:2115363-2115811**

TTTTTTGTAATATAAATGTATAGTCTTTCTCCTTTGTTTTCTCTCGTTCGTTTCCATGTTTCCAATTATGCTGTTCGTGTCATTCATTCTCCCTCCAATTTTCCCTTTTTGTGTGCGCCCTTCTAACTTCCCCTTCCCTCTAACGAGATATTTGCTTCGCTACGCTACGCACTCCCTACCATAATATACTCACTAACATTATACTTTCACTACATCTCATTACTTATTCCTACCATACACACTATATTTCAACACTTTCCTCACTCAACCATTCCATCAACCATCCTCTTTCTCCCCATACATCTTGAACTCCTTACACCCATACTTCATATCCATTACCCATTTTCCTTTATTAACATATATTAAACCTCATCCCAAATGTATCTACCCACTCTTTACATTCACATCCTTATTCCTTTAATTTCTTACGTTCCACTTCCTCAACCATA

**>JB4.chr2:2115363-2115811**

TTTTTTGTAATATAAATGTATAGTCTTTCTCCTTTGTTTTCTCTCGTTCGTTTCCATGTTTCCAATTATGCTGTTCGTGTCATTCATTCTCCCTCCAATTTTCCCTTTTTGTGTGCGCCCTTCTAACTTCCCCTTCCCTCTAACGAGATATTTGCTTCGCTACGCTACGCACTCCCTACCATAATATACTCACTAACATTATACTTTCACTACATCTCATTACTTATTCCTACCATACACACTATATTTCAACACTTTCCTCGCTCAACCACTCCATCAACCATCCTCTTTCTCCCCATACATCTTAAACTCCTTACACCCATACTTCATATCCATTACCCATTTTCCTTTATTAACATATATTAAACCTCATCCCAAATGTATCTACCCACTCTTTACATTCACATCCTTATTCCTTTAATTTCTTACCTTCCACTTCCTCAACCATA

**>JB594.chr2:2115363-2115811**

TTTTTTGTAATATAAATGTATAGTCTTTCTCCTTTGTTTTCTCTCGTTCGTTTCCATGTTTCCAATTATGCTGTTCGTGTCATTCATTCTCCCTCCAATTTTCCCTTTTTGTGTGCGCCCTTCTAACTTCCCCTTCCCTCTAACGAGATATTTGCTTCGCTACGCTACGCACTCCCTACCATAATATACTCACTAACATTATACTTTCACTACATCTCATTACTTATTCCTACCATACACACTATATTTCAACACTTTCCTCGCTCAACCACTCCATCAACCATCCTCTTTCTCCCCATACATCTTGAACTCCTTACACCAATACTTCATATCCATTAGCCATTTTCCTTTATTAACATATATTAAACCTCATCCCAAATGCATCTACCCACTCTTTACATTCACATCCTTATTCCTTTAATTTCTTACGTTCCACTTCCTCAACCATA

**>JB1171.chr2:2115363-2115811**

TTTTTTGTAATATAAATGTATAGTCTTTCTCCTTTGTTTTCTCTCGTTCGTTTCCATGTTTCCAATTATGCTGTTCGTGTCATTCATTCTCCCTCCAATTTTCCCTTTTTGTGTGCGCCCTTCTAACTTCCCCTTCCCTCTAACGAGATATTTGCTTCGCTACGCTACGCACTCCCTACCATAATATACTCACTAACATTATACTTTCACTACATCTCATTACTTATTCCTACCATACACACTATATTTCAACACTTTCCTCGCTCAACCACTCCATCAACCATCCTCTTTCTCCCCATACATCTTGAACTCCTTACACCAATACTTCATATCCATTAGCCATTTTCCTTTATTAACATATATTAAACCTCATCCCAAATGTATCTACCCACTCTTTACATTCACATCCTTATTCCTTTAATTTCTTACCTTCCACTTCCTCAACCATA

**>JB874.chr2:2115363-2115811**

TTTTTTGTAATATAAATGTATAGTCTTTCTCCTTTGTTTTCTCTCGTTCGTTTCCATGTTTCCAATTATGCTGTTCGTGTCATTCATTCTCCCTCCAATTTTCCCTTTTTGTGTGCGCCCTTCTAACTTCCCCTTCCCTCTAACGAGATATTTGCTTCGCTACGCTACGCACTCCCTACCATAATATACTCACTAACATTATACTTTCACTACATCTCATTACTTATTCCTACCATACACACTATATTTCAACACTTTCCTCGCTCAACCACTCCATCAACCATCCTCTTTCTCCCCATACATCTTGAACTCCTTACACCCATACTTCATATCCATTACCCATTTTCCTTTATTAACATATATTAAACCTCATCCCAAATGTATCTACCCACTCTTTACATTCACATCCTTATTCCTTTAATTTCTTACCTTCCACTTCCTCAACCATA

**>JB22;JB374;JB759;JB760;JB761;JB868;JB869;JB870;JB875;JB879;JB885;JB886;JB887;JB888;JB889;JB891;JB892;JB897;JB898;JB906;JB908;JB917;JB936;JB937;JB938;JB940;JB941;JB944;JB945;JB947;JB1111;JB1117;JB1168;JB1169;JB1170;JB1179;JB1181;JB1182;JB1183;JB1184;JB1195;JB1204.chr2:2115363-2115811**

TTTTTTGTAATATAAATGTATAGTCTTTCTCCTTTGTTTTCTCTCGTTCGTTTCCATGTTTCCAATTATGCTGTTCGTGTCATTCATTCTCCCTCCAATTTTCCCTTTTTGTGTGCGCCCTTCTACCTTCCCCTTCCCTCTAACGAGATATTTGCTTCGCTACGCTACGCACTCCCTACCATAATATACTCACTAATATTATACTTTCACTACATCTCATTACTCATTCCTACCATACACACTATACTTCAACACTTTCCTCGCTCAACCACTCCATCAACCATCCTCTTTCTCCCCATACATCTTGAACTCCTTACACCAATACTTCATATCCATTAGCCATTTTCCTTTATAAACATATATTAAACCTCATCCCAAATGCATCTACCCACTCTTTACATTCACATCCCTATTCCTTTAATATCTTACGTTCCACTTCCTCAACCATA

**>JB900;JB901;JB918;JB1114;JB1175;JB1189;JB1190;JB1191;JB1192;JB1193;JB1194.chr2:2115363-2115811**

TTTTTTGTAATATAAATGTATAGTCTTTCTCCTTTGTTTTCTCTCGTTCGTTTCCATGTTTCCAATTATGCTGTTCGTGTCATTCATTCTCCCTCCAATTTTCCCTTTTTGTGTGCGCCCTTCTACCTTCCCCTTCCCTCTAACGAGATATTTGCTTCGCTACGCTACGCACTCCCTACCATAATATACTCACTAATATTATACTTTCACTACATCTCATTACTCATTCCTACCATACACACTATACTTCAACACTTTCCTCGCTCAACCACTCCATCAACCATGCTCTTTCTCCCCATACATCTTGAACTCCTTACACCAATACTTCATATCCATTAGCCATTTTCCTTTATAAACATATATTAAACCTCATCCCAAATGCATCTACCCACTCTTTACATTCACATCCCTATTCCTTTAATATCTTACGTTCCACTTCCTCAACCATA

**>JB931.chr2:2115363-2115811**

TTTTTTGTAATATAAATGTATAGTCTTTCTCCTTTGTTTTCTCTCGTTCGTTTCCATGTTTCCAATTATGCTGTTCGTGTCATTCATTCTCCCTCCAATTTTCCCTTTTTGTGTGCGCCCTTCTACCTTCCCCTTCCCTCTAACGAGATATTTGCTTCGCTACGCTACGCACTCCCTACCATAATATACTCACTAATATTATACTTTCACTACATCTCATTACTCATTCCTACTATACACACTATACTTCAACACTTTCCTCGCTCAACCACTCCATCAACCATCCTCTTTCTCCCCATACATCTTGAACTCCTTACACCAATACTTCATATCCATTAGCCATTTTCCTTTATAAACATATATTAAACCTCATCCCAAATGCATCTACCCACTCTTTACATTCACATCCCTATTCCTTTAATATCTTACGTTCCACTTCCTCAACCATA

**>JB1197.chr2:2115363-2115811**

TTTTTTGTAATATAAATGTATAGTCTTTCTCCTTTGTTTTCTCTCGTTCGTTTCCATGTTTCCAATTATGCTGTTCGTGTCATTCTTTCTCCCTCCAATTTTCCCTTTTTGTGTGCGCCCTTCTAACTTCCCCTTCCCTCTAACGAGATATTTGCTTCGCTACGCTACGCACTCCCTACCATAATATACTCACTAACATTATACTTTCACTACATCTCATTACTCATTCCTACCATACACACTATATTTCAACACTTTCCTCACTCAACCATTCCATCAACCATCCTCTTTCTCACCATACATCTTGAACTCCTTACACCAATACTTCATATCCATTAGCCATTTTCCTTTATAAACATATATTAAACCTCATCCCAAATGCATCTACCCACTCTTTACATTCACATCCCTATTCCTTTAATATCTTACGTTCCACTTCCTCAACCATA

**>JB837.chr2:2115363-2115811**

TTTTTTGTAATATAAATGTATAGTCTTTCTCCTTTGTTTTCTCTCGTTCGTTTCCATGTTTCCCATTATGCTGTTCGCGTCATTCATTCTCCCTCCAATTTTCCCTTTTTGTGTGCGCCCTTCTAACTTCCCCTTCCCTCTAACGAGATATTTGCTTCGCTACGCTACGCACTCCCTACCATAATATACTCACTAACATTATACTTTCACTACATCTCATTACTTATTCCTACCATACACACTATATTTCAACACTTTCCTCGCTCAACCACTCCATCAACCATCCTCTTTCTCCCCATACATCTTGAACTCCTTACACCAATACTTCATATCCATTAGCCATTTTCCTTTATAAACATATATTAAACCTCATCCCAAATGCATCTACCCACTCTTTACATTCACATCCCTATTCCTTTAATATCTTACGTTCCACTTCCTCAACCATA
